# Machine learning approaches reveal genomic regions associated with sugarcane brown rust resistance

**DOI:** 10.1101/2020.03.10.985960

**Authors:** Alexandre Hild Aono, Estela Araujo Costa, Hugo Vianna Silva Rody, James Shiniti Nagai, Ricardo José Gonzaga Pimenta, Melina Cristina Mancini, Fernanda Raquel Camilo dos Santos, Luciana Rossini Pinto, Marcos Guimarães de Andrade Landell, Anete Pereira de Souza, Reginaldo Massanobu Kuroshu

## Abstract

Sugarcane is an economically important crop, but its genomic complexity has hindered advances in molecular approaches for genetic breeding. New cultivars are released based on the identification of interesting traits, and for sugarcane, brown rust resistance is a desirable characteristic due to the large economic impact of the disease. Although marker-assisted selection for rust resistance has been successful, the genes involved are still unknown, and the associated regions vary among cultivars, thus restricting methodological generalization. We used genotyping by sequencing of full-sib progeny to relate genomic regions with brown rust phenotypes. We established a pipeline to identify reliable SNPs in complex polyploid data, which were used for phenotypic prediction via machine learning. We identified 14,540 SNPs, which led to a mean prediction accuracy of 50% by using different models. We also tested feature selection algorithms to increase predictive accuracy, resulting in a reduced dataset with more explanatory power for rust phenotypes. Using different feature selection techniques, we achieved accuracy of up to 95% with a dataset of 131 SNPs related to brown rust QTL regions and auxiliary genes. Therefore, our novel strategy has the potential to assist studies of the genomic organization of brown rust resistance in sugarcane.

## Introduction

Sugarcane is an important source of income worldwide, especially due to its efficiency in the manufacturing of biofuel and sugar-related products in most tropical and subtropical areas of the world^1, 2^. Although this crop has great energetic potential, its breeding process has generated high genomic complexity across bred varieties, exceeding that of most if not all other crops^3^. Modern sugarcane cultivars are derived from a process of hybridization that has occurred over a century between *Saccharum spontaneum* (2*n* = 5*x* = 40 to 16*x* = 128; *x* = 8)^3, 4^ and *Saccharum officinarum* (2*n* = 8*x* = 80, *x* = 10)^3, 4^. *S. officinarum* has a more efficient process of sugar production but is susceptible to several biotic and abiotic stresses, in contrast to *S. spontaneum*, which has a low sucrose content but is resistant to different types of stress^1, 3, 5^. Sugarcane cultivars have unique chromosome sets (with numbers ranging from 80 to 130)^6^ with highly complex genomic organization^1^, a polyploid genome (with overall ploidy estimated to be between 6 and 14)^7^, a frequent occurrence of aneuploidy at the locus level depending on the number of homologous chromosomes in hybrid cultivars^8^, an estimated whole-genome size of 10 Gb^9^, and a high content of repetitive regions (50% of genome size)^10^. This complexity has challenged the efforts of the scientific community to unravel the genetic architecture of sugarcane in terms of the molecular mechanisms underlying different phenotypes, particularly efforts to detect regions of phenotype-genotype associations.

Sugarcane breeding programs are implemented with the intention of releasing new cultivars with interesting agronomic traits, including disease resistance^11^. One disease with a large impact on sugarcane yield is brown rust, which is caused by *Puccinia melanocephala*, a fungus that affects foliage and decreases the photosynthetic capacity of sugarcane^12, 13^. Brown rust infections have already caused large economic losses^14–16^. However, disease control has been shown to be successful in sugarcane breeding^17^, and the planting of cultivars resistant to brown rust is considered the most effective method of controlling this pathogen^11, 12^. Based on comparisons of the genetic characteristics of the resistant cultivar R570 and other sugarcane varieties^17^, brown rust resistance was found to be a dominant trait controlled by one or a few genes^11, 18^, with the presence of two related major genes: Bru1^19^ and Bru2^20^. Bru1 has already been employed in different breeding programs to identify resistant sugarcane genotypes^5^, using, for instance, the presence of flanking molecular markers for resistance diagnosis across cultivars^12^.

Although there have been several advances in understanding brown rust susceptibility in sugarcane, it is important to consider that pathogens may overcome the resistance of sugarcane varieties, and the use of a single region for resistance examination further increases the probability of vulnerability^5^. Therefore, the exploration of novel genes could contribute to the understanding of this process and in turn overcome the problems associated with reliance on a single gene^21^. An appropriate strategy for unraveling the genetic architecture and genomic organization of brown rust resistance would be the use of linkage maps followed by quantitative trait locus (QTL) identification. However, existing methodologies for the construction of saturated linkage maps with high resolution are not suitable for sugarcane and other autopolyploid species due to their genomic complexity^5, 22, 23^. Using simplification strategies based on the population expected segregation ratio, such as the selection of a subset of single-dose markers, leads to impaired linkage groups and thus compromises the identification of reliable QTLs^22^. A linkage map depicting QTLs associated with brown rust resistance has been published^5^, but as observed in previous studies^11, 24^, adjustments of existing methods resulted in gaps, a poorly saturated map and a large number of unlinked markers, mainly due to the high probability of meiotic behaviors in the cultivars and the aneuploidy of sugarcane^7, 22^. Different software programs have been developed to build linkage maps for polyploids^22, 25–27^; however, none address sugarcane genomic organization.

The use of Bru1 for marker-assisted selection (MAS) represents a successful application of this methodology in some sugarcane varieties^28^. However, resistance differs among cultivars, which can restrict the application of validated linked markers as a general tool for MAS^21, 28^. Therefore, the identification and characterization of brown rust resistance genes in sugarcane have been slow^14^, mainly because selection approaches based on QTL mapping overestimate the effect of strong QTLs, while weak QTLs might not be identified^29, 30^. In general, these methodologies have low power to detect rare variants with phenotypic associations^31^. Methodologies for addressing sugarcane genomic characteristics are still lacking, and because of the difficulty of accurately selecting QTL regions for MAS, an alternative methodology known as genomic selection (GS) has been developed to identify promising varieties with resistance traits and improve sugarcane breeding programs in terms of time and cost^32, 33^.

In general, GS is based on the creation of a predictive model for breeding values built with the entire set of markers using a training and a testing population. This model might be posteriorly applied in a breeding program to select a set of promising individuals^33^. In sugarcane breeding programs, the selection of superior genotypes might take more than 12 years^34^, and GS represents an alternative for improving this process, accelerating the breeding cycle and reducing the time needed to generate diversity^31, 33, 35^. Due to sugarcane’s genomic complexity, simplified predictive models involving linear regression cannot capture the unknown nonlinear characteristics present in these datasets^31^, as described for other polyploid species^36–38^. To address this issue, machine learning (ML) methodologies represent a promising approach with high accuracy^31, 39–41^. Although GS was developed to address the problem of categorizing individuals using different populations, its application in biparental populations is suitable and might be highly efficient due to the significant amount of linkage disequilibrium between loci^42^, which would facilitate the initial cycles of breeding programs.

In sugarcane, the allele dosages (ADs) of a locus are frequently unknown^7^, which might lead to misclassified genotypes. These difficulties in genotyping a population directly impact the estimation of locus effects on model creation^43^, and this influence is more complex when using nonlinear models with more parameters to be estimated^44^. An alternative for dealing with erroneous features and additional restrictions for high-dimensional data is feature selection (FS). These techniques aim to reduce the number of single nucleotide polymorphisms (SNPs) in a data set and identify a subset of markers with higher predictive capability by removing markers that are irrelevant/redundant for the phenotype^45^. These methods are among the most powerful alternatives for building better generalization models^46^ while avoiding overfitting and the attribution of nongenetic effects to different markers^43^. With FS, it is possible to reduce marker density and build simpler and more comprehensive models^46^, thereby increasing predictive power due to the identification of phenotype-associated polymorphisms. A few previous studies applied ML methods to decrease the number of SNP datasets needed for phenotypic predictions^47–49^, achieving high accuracy. The identification of such a subset of putative causal polymorphisms is crucial for improving production in plants^42^ and represents a novel strategy for genomic prediction in sugarcane.

Therefore, the objectives of this research were as follows: (1) genotyping a sugarcane full-sib population using a genotyping by sequencing (GBS) protocol^50^ followed by an established bioinformatics pipeline to identify reliable SNPs considering the sugarcane aneuploid condition; (2) creating a ML-based strategy to establish a subset of SNPs with good ability to predict brown rust phenotypes; and (3) examining these polymorphic regions to identify genes and QTL regions. Our study provides a novel methodology that can assist in sugarcane genetic studies and breeding programs to establish a pipeline to infer phenotype-causative regions, which can help unravel sugarcane brown rust resistance molecular mechanisms and identify targets for breeding.

## Material and Methods

### Mapping Population and Phenotypic Characterization

A set of full-sib progeny composed of 219 individuals derived from a biparental cross between the elite clone IACSP95-3018 (female parent) and the commercial variety IACSP93-3046 (male parent) was developed by the Sugarcane Breeding Program at the Agronomic Institute of Campinas (IAC). IACSP95-3018 is a promising clone that is used in breeding programs but is susceptible to brown rust. IACSP93-3046 is a variety with good tillering, an erect stool habit and resistance to brown rust. These parents have already been used in transcriptome^51^ and mapping studies^24, 52^.

The progeny phenotyped for brown rust symptoms were planted in 2005 at the Sugarcane Breeding Center of the Instituto Agronômico (IAC) located in Ribeirão Preto-SP, Brazil, and again in 2011 in Piracicaba-SP, Brazil, in an augmented block design with five blocks, each containing 44 individuals, plots with 1-m rows and plants spaced 1.5 m apart. Both parents and two varieties (SP81-3250 and RB835486) were included in each replicate as controls. The level of brown rust infection was evaluated using a diagrammatic scale between 1 and 9, with larger values indicating larger percentages of leaf area infection^53^. In Ribeirão Preto, four evaluations were performed: (1) November 2005 (plant cane), (2) January 2006 (plant cane), (3) January 2007 (ratoon cane), and (4) March 2007 (ratoon cane). In Piracicaba, the evaluations were conducted in December 2011 (plant cane) and in February 2012 (plant cane).

### Phenotypic Data Analyses

The phenotypic analyses of brown rust were performed using R statistical software^54^ following a statistical mixed model:

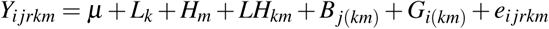

where *Y*_*i jrkm*_ is the phenotype of the *i*th genotype, considering the *j*th block, the *r*th replicate, the *k*th location and the *m*th year of harvest. The trait mean is represented by *µ*; the fixed effects were modeled to estimate the contributions of (1) the *k*th location (*L*_*k*_), (2) the *m*th harvest (*H*_*m*_), (3) the *j*th block at the *k*th location and in the *m*th harvest (*B*_*j*(*km*)_), and (4) the interaction between the *k*th location and *m*th harvest (*LH*_*km*_). The random effects included genotype *G* and the residual error *e*, representing nongenetic effects.

The residual distribution was evaluated using quantile-quantile (Q-Q) plots together with a Shapiro-Wilk normality test (p-value < 0.05). We also tested normalized values of the brown rust trait created with the R package bestNormalize^55^. To analyze the contribution of genotype to phenotype, we used best linear unbiased predictions (BLUPs) calculated based on the mixed model described above using the R package breedR^56^. Heritability (*H*^2^) was estimated as follows:

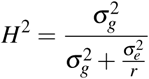

where 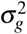 is the genetic variance, 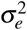 is the residual variance, and *r* is the number of replicates.

With these predictions, cluster analysis was performed with the BLUP values. We used complete hierarchical clustering based on pairwise Euclidean distances for visual inspection. The number of appropriate clusters was identified using the K-means algorithm together with (1) the within-cluster sums of squares and (2) the average silhouette width of clusters, implemented in the R package factoextra^57^. To evaluate the differences among the phenotypic rust groups, we used T-tests of the BLUPs and original values.

### Library Preparation and Sequencing Methodology

Total genomic DNA samples from parents and 180 progeny were extracted from leaf roll using the CTAB protocol^58^. Genome complexity was reduced via the *PstI* restriction enzyme for library preparation^50^. We constructed two 9,648-plex libraries from the population consisting of a single sample of each individual, two replicate samples of each parent and one blank sample. Five sequencing runs were performed with the Illumina GAIIx (one in 2015) and Illumina NextSeq (four in 2017 divided into two groups, which were sequenced twice) systems.

### Quality Filtering and Demultiplexing

PhiX sequences were removed from GBS reads through alignments of raw reads against the PhiX genome using BLASTn^59^. Reads resulting in a minimum percent identity of 90% and e-value of 0.01 against PhiX regions were filtered out^60^. FASTQC^61^ was used for the initial visualization of nucleotide distributions and their respective qualities, and FastX-Toolkit scripts^62^ were employed to obtain 90-bp reads with a minimum of 80% of bases with a Q greater than 20. Sample demultiplexing was also performed using the FastX-Toolkit^62^.

### Read Alignment and Reference Evaluation

We used the BWA-MEM version 0.7.12^63^ and Bowtie2 version 2.3.3.1^64^ algorithms to align the filtered reads against the following references: (1) the methyl-filtered (MF) genome of sugarcane cultivar SP70-1143^65^, (2) the sorghum genome from Phytozome v.13^66^, a sugarcane leaf transcriptome^51^, (4) the draft genome of the hybrid SP80-3280^67^, (5) the monoploid genome of the R570 variety^3^, (6) the *S. spontaneum* genome split into four subsets on the basis of the allele-defined genome, and (7) sequences from the sugarcane expressed sequence tag project (SUCEST)^68^.

The performance of each mapping software tool was evaluated according to the percentage of uniquely mapped reads. Additionally, to identify the most appropriate reference for SNP calling in sugarcane, SAMtools version 1.6^69^ was used to examine the profiles of sequencing depth across loci^69^.

Stacks version 2.3^70^ was used to obtain the consensus sequences (contigs) formed by mapping reads to the different references. All contigs were evaluated on the basis of their similarity to the consensus sequences of the other possible references. All correspondences were counted, taking into account the quantity of related contigs. These alignments were obtained through BLASTn^59^ with stringent parameters to examine real redundancies (a minimum e-value of 1e-30, a minimum percent identity of 95% and coverage of at least 75% in the query sequence). We also used the R package circlize^71^ to visually inspect these redundancies. Additionally, traditional assembly metrics (number of contigs, largest contig, total length, quantity of ambiguous bases (Ns) per 100 kbp, N50/75 and L50/L75) of all the contigs separated by reference were calculated using QUAST version 5.0.2^72^. An evaluation of the raw reference sequences was performed using QUAST version 5.0.2.

### SNP Calling and Ploidy Evaluation

With the different references used, we executed the Tassel4-POLY^73, 74^ and Stacks version 2.3^70^ pipelines. We evaluated raw SNPs by comparing them to a dataset with a filter criterion of a maximum of 25% of missing data per locus, considering individual genotypes without a minimum count of 50 reads as missing data.

The best selected reference was used to identify variants through the Haplotype Caller algorithm implemented in Genome Analysis Toolkit (GATK) version 3.7^75^, SAMtools version 1.6^69^ and FreeBayes version 1.1.0-3^76^. We created a common dataset to be processed by these tools, establishing a pre-processing pipeline according to GATK best practices^75^. From the mapping results, uniquely mapped reads were selected using SAMtools version 1.6^69^, and with Picard Toolkit^77^, the following steps were performed: (1) the mapped files from different sequencing experiments were joined into one file per individual; (2) read duplicates were marked; and (3) read group information was added to different files. To produce more accurate results, we used GATK version 3.7^75^ to realign indels and SAMtools version 1.6^69^ to convert mapping formats. Putative SNPs were called using the three different tools and different ploidy configurations with GATK and FreeBayes (even ploidies ranging from 2 to 20). We selected the identified SNPs and evaluated these variants with respect to the quantity of missing data.

### Final SNP-set Selection and Ploidy Evaluation

Using the R package VennDiagram^78^, a Venn diagram was created to evaluate the intersection between SNPs identified by the callers and those identified by the selected reference. Indels were not used for further analyses. Due to sugarcane aneuploidy at the locus level, we genotyped the individuals on the basis of SNP allele proportions, i.e., the ratio between the number of reads for the reference allele and the total number of reads. To increase the reliability of our results, we selected markers called by Tassel and at least one other caller with a minimum count of 50 reads per individual and a maximum of 25% missing data.

SuperMASSA^79^ and the VCF2SM pipeline^73^ were used to estimate the ploidy levels at different loci. Quantitative allele intensities at each locus were estimated for individuals based on read depth^73^. These values were used to estimate locus ploidies (ranging from 2 to 20). We used the F1 model for population structure due to the usage of a biparental population and did not restrict the posterior probability threshold to capture and analyze all possible configurations produced by the statistical estimate. We also defined the most probable set of loci with a posterior probability greater than 0.8 given the selected ploidy (6 through 14).

### Machine Learning Strategies

Using the identified SNPs as ADs and allele proportions (APs), eight ML algorithms were tested to check their ability to predict the phenotypic rust groups. Missing data were imputed as the means. We tested K-nearest neighbor (KNN)^80^, support vector machine (SVM)^81^, Gaussian process (GP)^82^, decision tree (DT)^83^, random forest (RF)^84^, multilayer perceptron (MLP) neural network^85^, adaptive boosting (AB)^86^, and Gaussian naive Bayes (GNB)^87^ implemented in the scikit-learn Python v.3 module^88^. As a cross-validation strategy, we used a stratified K-fold (k=4) repeated 100 times for different data configurations. We evaluated the following metrics: (1) accuracy (proportion of correctly classified items), (2) recall/sensitivity (items correctly classified as positive among the total quantity of positives), (3) precision (items correctly classified as positive among the total items identified as positive), and (4) specificity (items classified as negative among the total negative items). The area under the receiver operating characteristic (ROC) curve (AUC) was also calculated for each model and plotted using the Matplotlib library^89^ with Python v.3.

We also tested FS techniques implemented in the scikit-learn Python v.3 module^88^. We tested the following approaches to obtain feature importance and create subsets of the marker data: (1) gradient tree boosting (FS1)^90^, (2) L1-based FS through a linear support vector classification system (FS2)^81^, (3) extremely randomized trees (FS3)^91^, (4) univariate FS using ANOVA (FS4), and (5) RF (FS5)^84^. On the evaluation metrics (accuracy, recall, precision and specificity) for each subset of identified markers (FS1, FS2, FS3, FS4 and FS5), we performed a Shapiro-Wilk normality test (p-value < 0.01) and identified confidence intervals for means (95%, 99% and 99.9% confidence) using the gmodels R package^92^ for parametric data and a Wilcoxon test for nonparametric data. We tested the differences in these metrics between the selected FS methods using ANOVA and multiple comparisons by Tukey’s test implemented in the agricolae R package^93^. We also evaluated the intersection of these datasets using the R package VennDiagram^78^.

### Functional Annotation

From the SNPs identified by the most promising FS technique, we selected the reference positions to which they belonged and extracted the respective region. To check the distribution of these SNPs in the Bru1 region, we selected nine bacterial artificial chromosomes (BACs) from the sugarcane cultivar R570 that were previously described as belonging to regions containing Bru1^94^. These BACs were retrieved from the GenBank database^95^. We performed comparative alignments of the nine BACs and the selected reference sequences against *S. spontaneum*^1^ coding DNA sequences (CDSs) using BLASTn^59^ with the following parameters: a minimum e-value of 1e-30, a minimum percent identity of 95% and coverage of at least 75% in the query sequence. The distribution of these regions among *S. spontaneum* chromosomal regions was inferred using the karyoploteR package^96^.

We created a dataset with CDSs extracted from Phytozome v.13^66^ for fourteen different species from the Poaceae family (*Brachypodium distachyon, Brachypodium hybridum, Brachypodium silvatium, Hordeum vulgare, Oryza sativa, Oropetium thomaeum, Panicum hallii, Panicum virgatum, Sorghum bicolor, Setaria italica, Setaria viridis, Triticum aestivum, Thinopyrum intermedium* and *Zea mays*) and *Arabidopsis thaliana*. Selected *S. spontaneum* CDSs were aligned against this dataset, enabling the identification of correspondence with Gene Ontology^97^ (GO) categories and Kyoto Encyclopedia of Genes and Genomes (KEGG) Orthologies^98^ (KOs). All identified GO categories were used to create a treemap to visualize possible correlated categories in the dataset caused by identified regions using FS and BAC correspondence. This step was performed using the REVIGO tool^99^.

## Results

### Phenotypic Analyses

The brown rust phenotypic dataset was analyzed as described in Section 2 (Supplementary Figs. S1-S6) of the Supplementary Information (SI). Using the created phenotypic mixed model, we obtained a heritability of approximately 77%. Through the established statistical analysis procedures, we identified two different phenotypic groups, which were used for association analyses. These groups presented high divergence in scores, and the individuals were classified as belonging to the “resistant” group or the “susceptible” group.

### Genotyping Process

Raw GBS data quality control (Supplementary Tables S1 and S2), read alignment (Supplementary Table S3 and Supplementary Fig. S7), reference evaluation (Supplementary Tables S4-S9 and Supplementary Fig. S8) and SNP calling (Supplementary Tables S8-S11) were performed as described in the SI. The selected mapping tool was BWA, as it allowed the identification of a larger quantity of uniquely mapped reads. We considered the MF genome the most appropriate reference for SNP calling with our GBS dataset; this choice was made because this reference provided the largest percentage of uniquely mapped reads (Supplementary Table S3), the most consensus sequences and respective profiles (Supplementary Tables S4 and S5), the greatest sequencing depth at different mapping positions (Supplementary Table S6), the greatest ability for its consensus contigs to represent the majority of the other reference consensus sequences (Supplementary Table S7), and the largest quantity of SNPs identified using Tassel and Stacks (Supplementary Tables S8 and S9). The SNP calling process performed with the different tools and MF reference resulted in different quantities of markers, as described in the SI. The quantity of SNPs can be observed in Table 1.

**Table 1.** Final SNP sets obtained using the MF reference and different tools.

| Quantity | Raw SNPs | Biallelic SNPs |
| --- | --- | --- |
| Tassel | 135,979 | 135,594 |
| Stacks | 106,881 | 106,881 |
| GATK | 61,023 | 60,338 |
| SAMtools | 353,715 | 349,574 |
| FreeBayes | 72,391 | 71,999 |

The intersections between SNPs found with different tools can be visualized in Fig. 1. A total of 13,458 SNP markers were found by all used callers. However, when applying a reasonable filter for locus depths (minimum count of 50 per individual) and missing data (maximum of 25%)^23^, this quantity decreased to 2,284. Although this approach of selecting intersecting SNPs enables the definition of a highly stringent set, the quantity of false negatives will also be high. To establish a reasonable approach for SNP identification in sugarcane, we selected the most probable SNPs as the variants found with Tassel and at least one other caller. With this approach, we found 88,395 SNPs (eliminating 49,362 possibly false-positive SNPs uniquely identified by Tassel). After applying filters based on missing data and read depth, we obtained a final set of 14,540 SNPs (eliminating 4,341 questionable SNPs among 18,881 markers that would have been obtained using Tassel as a unique tool together with the described filters). These datasets were used to evaluate possible ploidy configurations using SuperMASSA software. This evaluation was performed on (I) 14,540 SNPs representing our final set of markers and (II) 4,341 SNPs representing the most likely false-positive SNPs.

**Figure 1.**
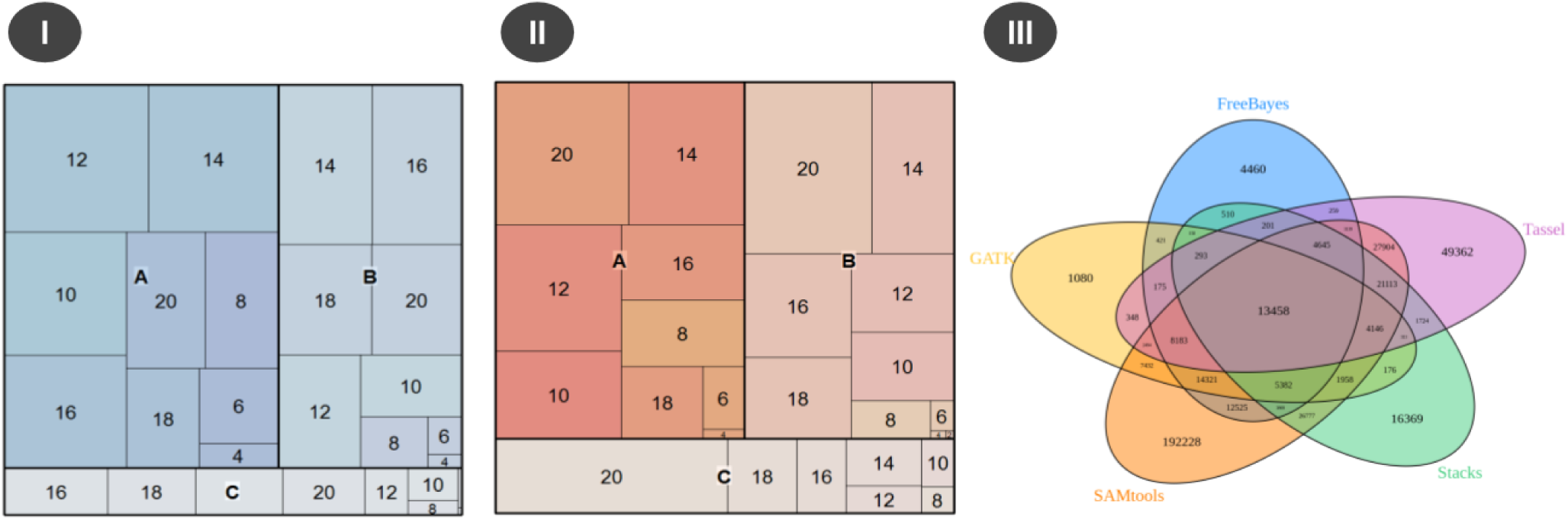
Figures (I) and (II) present ploidy estimates based on the intersection of SNPs between Tassel and at least one other tool (I) and Tassel-filtered SNPs not present in the intersection (II). Categories A, B and C are based on posterior probabilities (A represents probabilities larger than or equal to 0.8, B represents probabilities between 0.5 and 0.8, and C represents probabilities smaller than 0.5). In (III), a Venn diagram of SNPs called using the MF reference together with GATK, SAMtools, FreeBayes, Tassel and Stacks is shown.

Separating SuperMASSA posterior probabilities into three categories (A, B and C) based on their reliability (Fig. 1) classified a considerable number of SNPs as having a specific ploidy with high confidence. However, the first set with 14,540 SNPs included variants with ploidies more similar to those expected for sugarcane (6 to 14)^7^ compared with the second one with 4,341 SNPs. The majority of SNPs in the second set were classified as having a ploidy of 20, representing doubtful regions with chances of duplication events and low-quality data^7^. Therefore, this group of putative molecular markers with higher reliability provided better results in terms of estimated ploidies.

### Phenotype-Genotype Associations

To understand the genotypic associations with different brown rust phenotypes more generally, we chose to perform genotype-phenotype analyses with the phenotypic rust groups identified in the clustering analysis. We performed these tests using two different approaches for genomic prediction: ADs obtained with SuperMASSA (ploidy range between 6 and 14 and posterior probability greater than or equal to 0.8) and APs calculated based on Tassel output for read counts. With these two different datasets, the FS techniques were applied and generated different sets of SNPs (Table 2).

**Table 2.** Quantity of SNPs selected by the FS techniques when using allele proportions (APs) and allele dosages (ADs).

| Method | SNPs (AD) | SNPs (AP) |
| --- | --- | --- |
| Total SNP dataset (All) | 5,224 | 14,540 |
| Gradient Tree Boosting (FS1) | 283 | 345 |
| L1-based SVC (FS2) | 135 | 140 |
| Extra Trees (FS3) | 1,991 | 2,595 |
| F statistic from ANOVA (FS4) | 327 | 922 |
| Random Forests (FS5) | 1,345 | 1,253 |

These SNPs were used to predict the phenotypic rust groups using the eight selected ML algorithms in the proposed cross-validation scenario (described in SI in Supplementary Tables S12-S17). The performance of APs was superior to that of ADs for all evaluated metrics (Supplementary Table S18). In almost 73% of the tests with different algorithms, the usage of APs was equal or superior to that of ADs. Although there were some discrepancies across models and between FS subsets, we observed better use of sugarcane GBS data with APs. In addition, the quantity of SNPs discarded to obtain favorable ADs in sugarcane was almost 64%. Therefore, we considered the analysis of APs better than ADs for the task of GP.

The capability of predicting brown rust phenotypic groups was quite different among the created scenarios. Using the entire dataset, the overall accuracy was near 50%, showing the models’ inefficiency in capturing the real SNP effects. The KNN model presented an accuracy of almost 70% but with a very small value of specificity (0.23%), thus proving its inefficiency in predicting these phenotypes when using the entire set of SNPs. With FS techniques, these values increased but still presented differences between the selected methods. To evaluate the best FS techniques with which to increase predictive capabilities, we determined confidence intervals for all metric means (accuracy, recall, precision and specificity), as shown in Supplementary Table S19. Then, we counted the quantity of measures that exceeded the superior boundaries. FS1, FS2 and FS4 had the best performance, as described in the SI (Supplementary Tables S20-S23). Furthermore, we analyzed the distributions and similarities of these metrics. The accuracy distribution is shown in Fig. 2, and the other distributions are shown in the SI (Supplementary Figs. S9-S12). FS3 and FS5 clearly did not allow a substantial increase in these performance measures. The maximum values in the boxplots for FS3 and FS5 are close to the medians of FS1, FS2 and FS4. In addition, considering that multiple comparisons by Tukey’s test grouped F3, F5 and the initial dataset together, we can conclude that these techniques did not enable substantial improvement in accuracy. Analyses of the other metrics also showed better performance of FS1, FS2 and FS4 than of the other datasets, including FS3, FS5 and the entire set of SNPs. Due to these findings, we considered FS1, FS2 and FS4 the most promising methodologies for detecting variants with high predictive capabilities.

**Figure 2.**
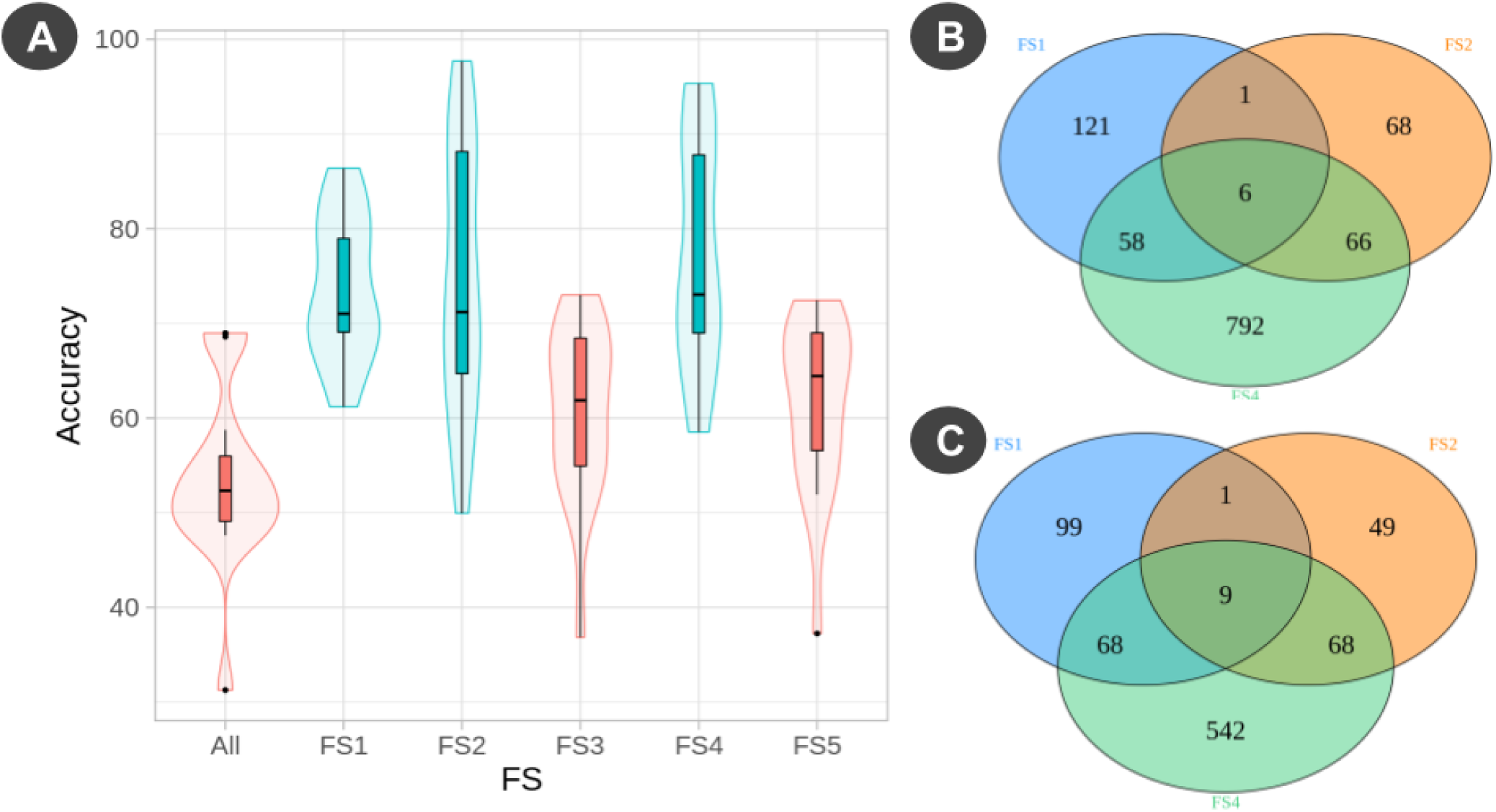
(**a**) Accuracy distribution using the different datasets for the ML strategies separated by FS strategy and colored based on the groups identified with multiple comparisons by Tukey’s test (ANOVA p-value of 0.000000000000455); (**b**) intersection of SNPs selected from FS1, FS2 and FS4; (**c**) intersection of MF scaffolds selected from FS1, FS2 and FS4.

The FS1, FS2 and FS4 methods identified different variants in different scaffolds. However, there were intersections between these sets (Fig. 2), which we decided to evaluate. We tested all selected ML algorithms using the intersection between at least two strategies (Inter 2), which corresponded to 131 SNPs, and the intersection between the three strategies (Inter 3), which corresponded to 6 SNPs. The results obtained using Inter 3 did not increase the metric values of the FS techniques; however, they were far superior to the initial results obtained with the entire dataset (approximately 41% larger), as described in the SI (Supplementary Table S24). Inter 2, however, showed the highest predictive capabilities (Table 3), suggesting that these variants have a greater probability of being associated with brown rust phenotypes.

**Table 3.** Performance of the intersection of FS1, FS2 and FS4 compared with the other strategies.

| Performance |  |  |  |  |
| --- | --- | --- | --- | --- |
| Model | Accuracy (AP) | Recall (AP) | Precision (AP) | Specificity (AP) |
| AB | 79.70 | 80.71 | 88.84 | 77.43 |
| DT | 64.71 | 64.84 | 80.23 | 64.43 |
| GP | 91.54 | 93.18 | 94.49 | 87.91 |
| KNN | 77.61 | 98.43 | 76.12 | 31.26 |
| MLP | 94.93 | 94.53 | 98.05 | 95.81 |
| GNB | 92.04 | 91.17 | 97.12 | 93.98 |
| RF | 72.29 | 67.15 | 90.19 | 83.74 |
| SVM | 87.29 | 84.44 | 96.73 | 93.64 |
| Means |  |  |  |  |
| Mean All | 51.14 | 47.77 | 71.37 | 57.24 |
| Mean FS1 | 77.80 | 80.54 | 86.89 | 72.94 |
| Mean FS2 | 82.01 | **89.60** | 86.70 | 76.69 |
| Mean FS4 | 77.61 | 86.67 | 85.80 | 74.75 |
| Mean Inter 2 | **83.50** | 87.81 | **92.34** | **85.83** |
| Mean Inter 3 | 72.28 | 72.77 | 84.90 | 71.18 |

The tested ML models had different capabilities of separating the phenotypic groups, and these capabilities changed depending on the dataset used. In addition to using the previous metrics, we chose to evaluate model performance using ROC curves and the respective AUCs. All of these plots are shown in the SI together with the AUC values (Supplementary Figs. S13-S20). We evaluated two different configurations to consider a model with reasonable predictive performance: (A) AUC *≥* 0.8 and (B) AUC *≥* 0.9%. For (A), we identified AB, GNB, GP and MLP as the most promising models when using the FS1, FS2 and FS4 techniques. When using the entire dataset and FS3, there were no significant changes in performance under (A). GNB was the best model for FS5; GNB and RF were the best models for Inter 3; and KNN, GP, RF, MLP, AB and GNB were the best models for Inter 2. This first configuration enabled identification of the Inter 2 FS technique as the most appropriate for the creation of stable models using ML strategies. The performance of the built models based on Inter 2 is shown by ROC curves in Fig. 3 and contrasted with the results for the entire dataset. For (B), the entire dataset, FS3, FS5 and Inter 3 did not have AUC values exceeding 0.9, supporting the exclusion of FS3 and FS5 as interesting for detecting phenotype-associated variants. GNB was the best model for FS1, and GP, MLP and GNB were the best models for FS2, FS4 and Inter 2. Thus, we considered GP, MLP and GNB the best models for predicting the brown rust phenotypic groups. The ROC curves for these three algorithms and the different subsets are provided in the SI (Supplementary Figs. S21-S23). The best AUC values were (I) MLP: 0.99 for Inter 2 and 1.00 for FS2, (II) GNB: 0.98 for Inter 2 and 0.96 for FS2, and (III) GP: 0.98 for Inter 2 and 0.98 for FS2. This finding supports the hypothesis of an association between Inter 2 regions and brown rust phenotypes. On the basis of these results, we suggest that the identification of intersections between FS1, FS2 and FS3 might be an appropriate methodology for both GP and the identification of regions associated with brown rust phenotypes.

**Figure 3.**
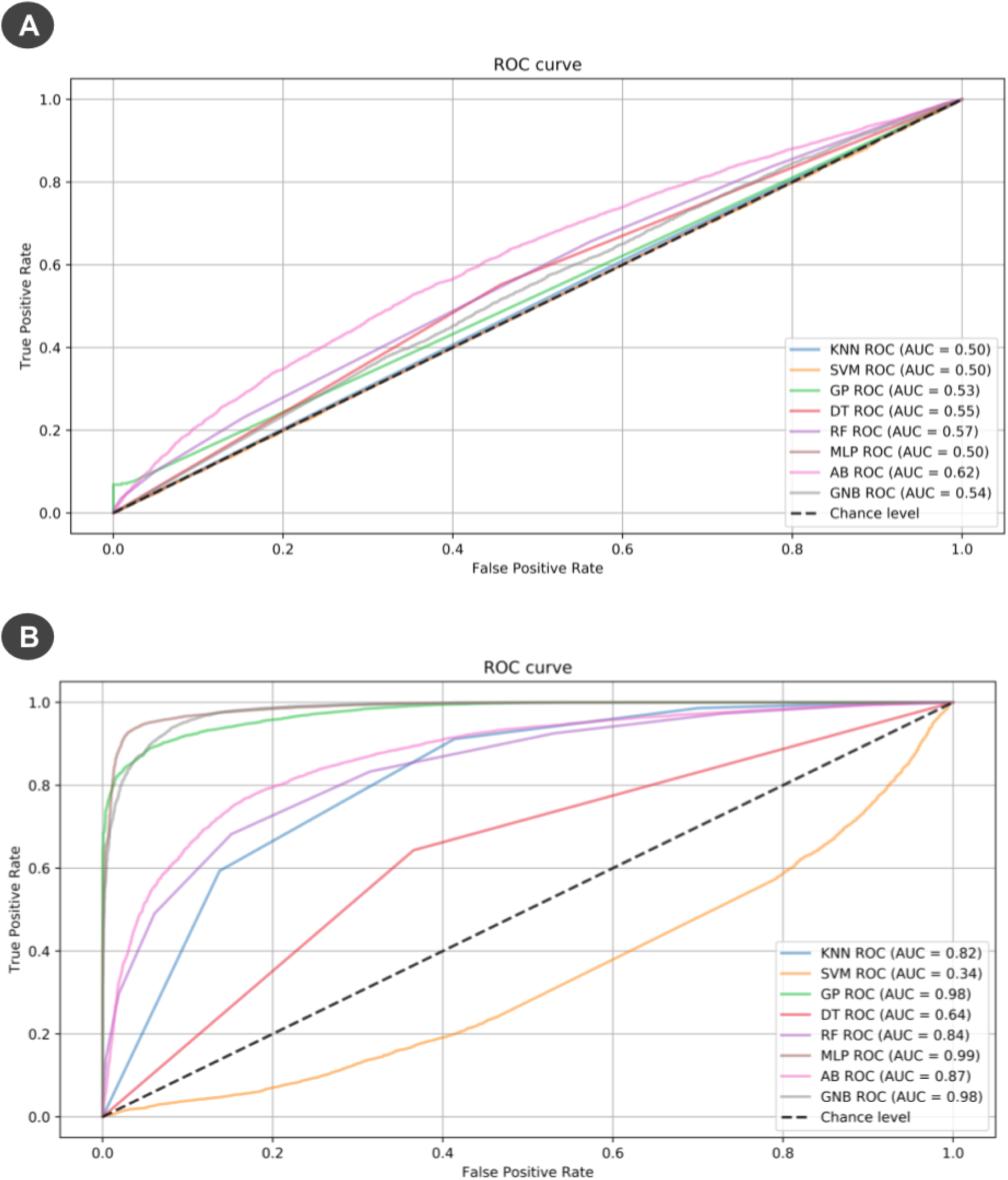
Receiving operating characteristic (ROC) curves showing the performance of the 8 ML strategies when using (**a**) the entire dataset of SNPs (14,540 SNPs) or (**b**) the SNPs shared by FS1, FS2 and FS4 (131 SNPs).

The last analysis that we performed to test whether this methodology was a promising strategy was an evaluation of the genomic regions where the selected variants were located. For this step, we used *S. spontaneum* CDSs corresponding to (A) 9 selected BACs related to Bru1 QTL regions and (B) 146 MF scaffolds identified as important by at least two methods. We identified 373 CDSs using (A) and 240 CDSs using (B). All BACs of (A) had correspondences, and nine scaffolds of (B) did not have relevant alignments. As there was only one CDS in common between (A) and (B), we evaluated the chromosomal location of these CDSs considering the *S. spontaneum* genomic reference, which is presented in the SI (Supplementary Fig. 24). Notably, regions where these CDSs were located were spread throughout the genome. However, nearly all CDSs identified in (B) were close to CDSs identified in (A), suggesting linkage disequilibrium between these regions due to chromosomal proximity. Additionally, to understand whether these genomic regions have similar impacts on biological processes, we performed enrichment analysis using the GO categories of these two groups (Fig. 4). We found 148 different GO categories in (A) and 100 in (B), with 50 GOs in common. The other 50 categories identified for only the selected variants can be found in the SI (Supplementary Fig. S25); there were four main categories: (I) sphingolipid metabolism, (II) DNA topological change, nitrogen compound transport, and (IV) phosphatidylinositol-mediated signaling.

**Figure 4.**
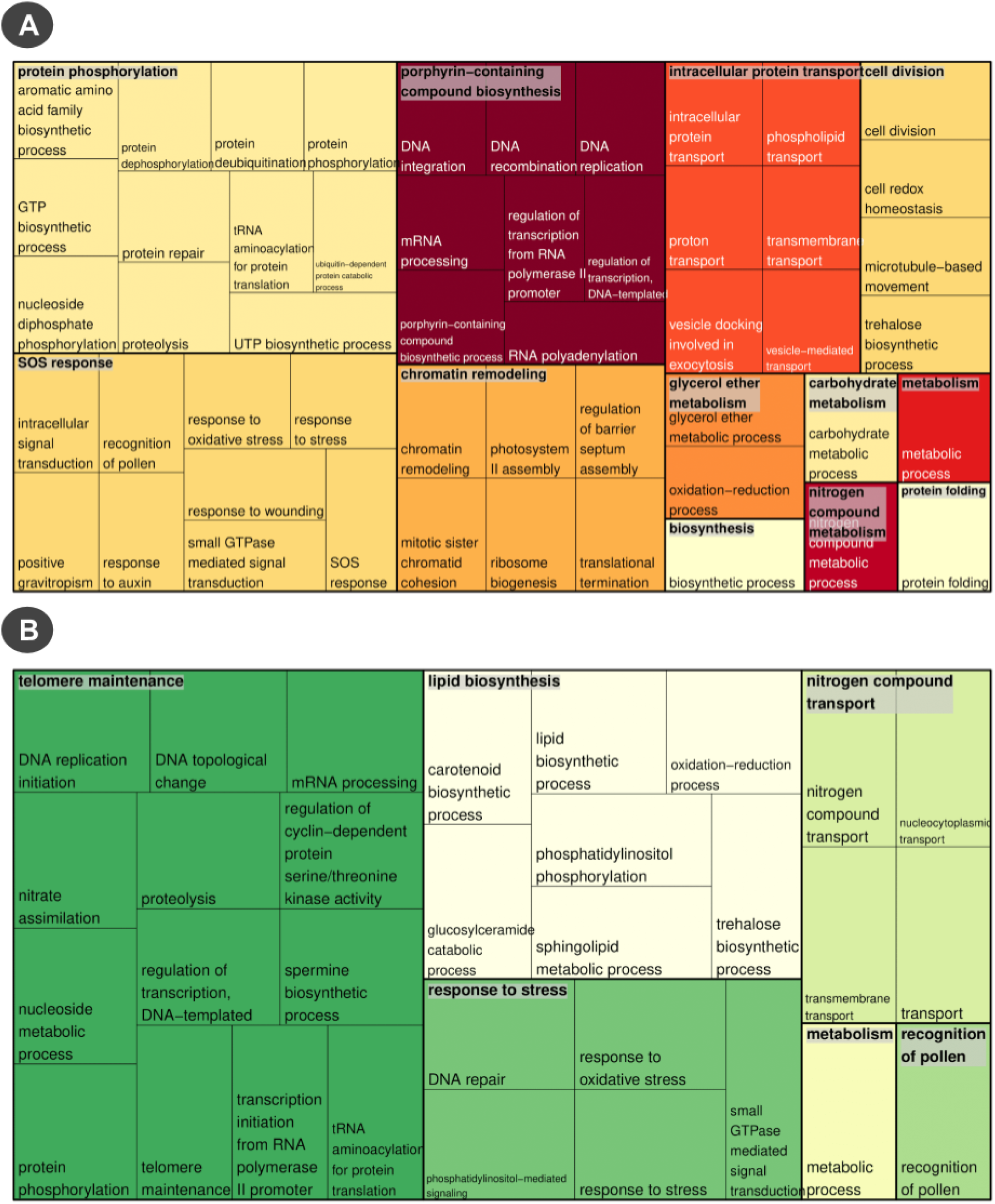
Treemaps plotted based on GO categories identified in (**a**) *S. spontaneum* CDSs corresponding to BAC sequences related to brown rust QTLs and (**b**) *S. spontaneum* CDSs corresponding to the scaffolds with the SNPs identified from the Inter 2 dataset.

In relation to metabolic pathways, we selected the *S. bicolor* KEGG correspondences for each CDS and also separated these findings into groups (A) and (B). The complete discrimination of the identified pathways is shown in the SI. We found 41 associated pathways in (A) and 29 in (B), with 16 in common between these groups. As expected, there was an elevated number of common biological cascades that might be influenced by these regions. The specific pathways found exclusively in group (B) were monoterpenoid biosynthesis, phenylpropanoid biosynthesis, the pentose phosphate pathway, sulfur metabolism, other glycan degradation, fatty acid elongation, basal transcription factors, ubiquitin-mediated proteolysis, various types of N-glycan biosynthesis, tryptophan metabolism, sphingolipid metabolism, carbon metabolism and N-glycan biosynthesis.

## Discussion

The organization of the sugarcane genome greatly challenges genetic studies of this species, and alternative approaches must be employed to overcome these difficulties. Here, we developed a novel strategy to address sugarcane genomic specificities and enable the identification of genomic regions related to brown rust resistance through the evaluation of ML predictive performance. The sequencing method, SNP detection process and phenotypic associations were designed to fit these singularities. Sugarcane brown rust susceptibility was previously studied and applied in sugarcane breeding programs^28^; however, there is still a gap in the characterization of the wide range of genes involved in the process of infection and how different genomic polymorphisms can influence this phenotype. The adjustments performed on these analyses showed reasonable results, and the identification of these possibly phenotype-causative regions can help unravel sugarcane brown rust resistance molecular mechanisms and the selection of targets for breeding.

First, due to the diversity of rust scores (1-9), the variation in rust phenotypes within populations and the qualitative nature of rust phenotypes, we decided to use the two groups identified by the BLUP clustering analysis instead of the raw scores. We were interested in finding markers and genomic regions related to brown rust resistance, and the establishment of these two major groups enabled the identification of resistance categories in the population. As previously described, these phenotypic rust groups presented a high level of differentiation in rust scores, and this contrast in susceptibility may aid in the identification of the most promising plants for sugarcane breeding programs. In addition, the establishment of these groups allowed the use of a wide range of ML strategies.

In relation to the sugarcane genotyping process, different approaches have been adopted by the scientific community to reduce the genomic complexity of sugarcane and utilize a limited amount of information. Song et al.^100^, for example, designed different probes using in silico approaches. The resulting regions were posteriorly adopted in other studies^101–104^ due to the large quantity of markers with sufficient sequencing depth located in genic regions. Another approach is GBS^5, 23, 28, 105–107^, which is the preferred genotyping method for plants with some degree of genomic complexity^23, 108^ mainly due to its simplicity, reproducibility and considerable genome coverage^109^. In addition, regulatory regions controlling different phenotypes are often located in noncoding DNA, and GBS allows the amplification of such regions^50^. Herein, we decided to use GBS to obtain a broader set of genomic regions with their respective probabilities of correspondence with rust resistance.

Sequencing reads are generally organized by using the *S. bicolor* genome for comparative alignments and the subsequent identification of putative variants with bioinformatic methods. This reference choice is due to sorghum’s phylogenetic proximity to sugarcane^5, 28, 105–107^ and, in some cases, probe experimental design^100–104^. Despite sorghum’s genome usage, the availability of sugarcane pseudoreferences has provided new genomic tools for scientific research as initially explored by Balsalobre et al.^23^, who used the sorghum genome, a sugarcane MF genome^65^, a sugarcane leaf transcriptome^51^ and SUCEST tags^68^. However, new sugarcane genomic resources are now available, such as the draft genome of the cultivar SP80-3280^67^, the monoploid genome of the R570 variety^3^ and the genome of the AP85-441 *S. spontaneum* cultivar^1^, which are phylogenetically closer to current sugarcane cultivar resources than are sorghum resources. Therefore, there is a need to explore these new references and check their appropriateness. As there are no previous reports of the usage of these novel references together with sugarcane GBS data, we decided to test them in order to identify the most appropriate reference.

Although GBS allows a reduction in genomic complexity, we must consider sugarcane singularities to establish an analysis pipeline. In GBS experiments, the consensus of read clusters at cutting sites could be adopted as a reference in cases where there is no appropriate sequence to use^50^. However, genome assembly is a difficult task when dealing with repetitive regions and polyploids^110^. With the aim of reducing possible biases, we decided not to use de novo approaches, which were previously described as inappropriate for sugarcane GBS data^105^.

In our study, the combination of BWA and MF scaffolds had the best performance for GBS data. BWA was previously reported as a sensitive tool for aligning sugarcane reads and retaining a large number of uniquely mapped sequences^100^. In terms of MF performance, this may be explained by the experimental procedures of MF sequencing and GBS library preparation. GBS library construction is based on the selection of a subset of genomic regions using methylation-sensitive restriction enzymes, which avoid repetitive regions^50^. To select our GBS regions, we used the enzyme PstI, which is a methyl-sensitive restriction enzyme, to select hypomethylated DNA^111^. Similarly, the MF genome was obtained through a process of sequencing where genomic regions were also selected based on hypomethylation^65^. This approach generated high compatibility between our data and the genomic reference, as observed in the comparative alignments and previous reports^23^. Although there have been great advances in understanding the sugarcane genome since the *S. spontaneum* genome became available, we decided to perform our analyses using the sugarcane MF genome to capture the most probable markers and establish a criterion based on data appropriateness. This genomic reference is still at the scaffold level, but as shown in this study, there is a high rate of redundancy among consensus sequences obtained through GBS data alignments with the different references. Due to this observed redundancy, we chose not to use all of the references. In addition to adding redundant markers, it is important to note that these different consensus contigs built based on different references can lead to different alignments of GBS data. These alignments may in turn produce different organizational profiles of read alignments and divergent SNPs. Therefore, we selected the most reference with the best usage of the amount of GBS data as the most appropriate and analyzed the respective SNPs.

A wide range of SNP callers are available. Tassel was developed to handle GBS data and has been widely applied to species with different genomic organizations. Although this tool enables the identification of many SNPs, it was previously described as insufficiently accurate to be used alone^112^. Thus, to increase the reliability of our data, we decided to use other SNP callers (GATK, FreeBayes, SAMtools and Stacks) in combination with Tassel, as the usage of SNPs identified by more than one caller is more reliable than the usage of SNPs identified by only one caller^113^. The intersection between the SNPs identified by at least two tools was established to increase the accuracy of these variants without substantially increasing the number of false negatives. In addition, Tassel was used due to its targeted development for GBS data and preprocessing steps. The Tassel workflow keeps read depths unchanged between the initial mapping and the final data generated for the identified genotypes. In sugarcane, this information is necessary to estimate ADs or calculate APs. Using this intersection approach, we identified the final set of SNPs to be used for our association analyses. Indels, however, were not selected. These variants identified by in silico strategies do not provide reliable information, showing elevated divergence between the existent callers and a probability of producing spurious variants^114^.

Using this approach, we found 14,540 putative SNPs. With these regions, we tested two different strategies for genotyping the population at these loci: (1) the usage of ADs estimated with SuperMASSA and (2) the usage of APs calculated based on Tassel output. For SuperMASSA estimations, we kept only SNPs with an estimated ploidy between 6 and 14 (minimum posterior probability of 0.8) due to sugarcane genomic configurations^7, 23^. However, sugarcane aneuploidy together with the common occurrence of duplication events might have influenced the process of estimating locus ploidies and, in turn, the process of categorizing the related dosages through the established filters. In addition, 64% of the identified SNPs were discarded when using this approach for obtaining dosages. Because we would not need to calculate chromosomal distances between loci for linkage map construction, the elevated loss of markers and the reduced performance of ADs in the task of genomic prediction, we decided to continue our analyses with APs. Previous tests of this approach yielded reasonable results^101, 102, 104^.

After establishing the bioinformatics pipeline for identifying and evaluating these regions, we studied the influence of SNP subsets identified by FS techniques on the task of predicting phenotypic rust groups. The amount of data generated by high-throughput sequencing technologies^115^ represents a challenge in genomic prediction, particularly due to the difficulty of working with high-dimensional datasets, i.e., the ‘large p, small n’ problem^116^. This increase in the amount of available information makes the task of directly applying these marker data in genomic analyses more difficult and necessitates appropriate preprocessing steps^117^. In this study, we proposed the use of FS techniques to select a smaller set of SNPs with more predictive power than the entire dataset and closer associations with the brown rust phenotype to assist the identification of regions associated with disease status. This can be considered quite advantageous in the context of genomic selection because the identification of a subset of markers allows a reduction in sequencing costs^49^. In addition, it has already been demonstrated that for genomic selection, a selected reduced number of SNPs has reasonable reliability^49, 118, 119^.

The identification of markers related to this phenotype using FS is based on these techniques to provide an interpretable model due to the close relation between trait and genotype; i.e., using the subset of high-density markers might help elucidate the regions most likely to be involved in phenotypic differentiation^120^. This strategy of selecting a subgroup of SNPs with higher predictive power and closeness to the predictive class has already been employed in different contexts^48, 121, 122^. In this study, we tested five different strategies and found three promising alternatives for executing this methodology. FS1, FS2 and FS4 substantially increased the models’ capabilities of predicting the phenotypic groups as demonstrated in this paper. We believe that this increase in predictive power is due to the identification of regions influencing the phenotype, possibly in QTLs or regulatory genomic elements. As a final strategy for the prediction and selection of these associated regions, we suggest the use of the intersection of these three techniques. This approach enabled the creation of more stable models using different ML algorithms and better accuracies for predicting these phenotypes.

Corroborating this hypothesis, we also found that most of the identified regions containing these SNPs were associated with QTLs with known biological functions, and there were also additional categories known to be correlated with rust resistance. Through comparative alignments between MF scaffolds and *S. spontaneum* CDSs, we identified these regions and compared them with CDSs correlated with BACs developed based on Bru1 regions. A total of 146 different scaffolds were selected as important for this predictive task by at least two methods (FS1, FS2 and FS4). Among these sequences, only 9 did not have correspondence with *S. spontaneum* CDSs, possibly due to the presence of additional noncoding regulatory elements. These regions can be targets of genetic studies due to their relationships with predicted phenotypes. Although there was no considerable intersection between CDSs associated with BACs and the selected scaffolds, we did find consensus in correlated biological functions. This divergence between regions is mainly explained by the differences between the populations used to generate the GBS data and the brown rust QTLs (which were used to select BACs). QTL regions are identified for a specific population, and there might be differences between datasets from different populations, especially for the sugarcane genome. In addition, the creation of sugarcane linkage maps relies on many adaptations of methods, such as the selection of only single-dosage markers^23^, which might lead to the identification of a restricted set of QTLs and the nonuse of many auxiliary genomic elements.

The exclusive GO categories related to the selected variants have already been reported to be associated with resistance. Sphingolipid metabolism is intimately connected to programmed cell death^123–125^; DNA topological change is a wider category with different implications in many biological processes, including responses to pathogens^126^; differences in nitrogen compound transport might be related to the accumulation of this nutrient and its influence on resistance against pathogens^127^; and phosphatidylinositol-mediated signaling includes important categories that also act on plants’ responses to pathogens^125^. A considerable number of metabolic pathways related to both BACs and the selected scaffolds were also detected. However, specific pathways were found to be associated with these scaffolds, mainly due to the different roles of the proteins encoded by these identified CDSs and because these pathways were already reported as being associated with plant responses to different pathogens^123, 128–135^, further corroborating our findings. The indication of possible mutation events in these regions provides evidence of differences in protein expression and phenotypic characteristics.

The identified regions with putative variants and high predictive performance for brown rust phenotypic groups can be employed as novel regions to investigate susceptibility-related traits. This proposed strategy can complement traditional methodologies for deciphering sugarcane genomic regions associated with pathogen infection responses and susceptibility. Although these SNPs were identified for only one biparental population, the strategy can be used for different populations, and the genes can be further investigated to validate the influence of the genomic regions on different phenotypes. This study represents an initial step in employing ML and FS strategies in sugarcane genomic studies. We illustrated the great potential of applying these methodologies to predict phenotypes by using a highly complex polyploid species.

## Acknowledgments

This work was supported by grants from the Fundação de Amparo à Pesquisa do Estado de São Paulo (FAPESP 2008/52197-4 and 2005/55258-6) and Coordenação de Aperfeiçoamento de Pessoal de Nível Superior (CAPES, Computational Biology Programme). A.A. received a PhD fellowship from FAPESP (2019/03232-6); E.C. received a PhD fellowship from FAPESP (2010/50031-1); R.P. received a MSc fellowship from FAPESP (2018/18588-8); and M.M. received a PD fellowship from FAPESP (2014/11482-9) and CAPES (88882.160095/2013-01).

## Author contribution statement

A.A. performed all analyses and wrote the manuscript; E.C. created the GBS library and performed the sequencing experiments; H.R. assisted in the execution of GBS quality control procedures and preprocessing steps; J.N. collaborated in the creation of ML models; R.P. and M.M. contributed to manuscript writing; F.S., L.P. and M.L. were responsible for the phenotypic experiments; and A.S. and R.K. conceived the project. All authors reviewed, read and approved the manuscript.

## Accession Codes

Sequencing data are available through the Sequence Read Archive (SRA) database with the accession code SRP151376.

