## Supplementary Information for "Machine learning approaches reveal genomic regions associated with sugarcane brown rust resistance"

### 1. Population Organization

The biparental population included: (1) the parents IACSP95-3018 and IACSP93-3046, named 3018 and 3046, respectively; (2) the controls SP81-3250 and RB835486, named 3250 and 5486, respectively; and (3) the progeny composed of 219 individuals, numbered from 1 to 220 (with the exception of 57, which died during the experiment). Genotypic data were obtained for 180 individuals, named: 3, 5, 7, 8, 13, 14, 15, 18, 20, 21, 23, 24, 25, 26, 27, 29, 30, 33, 34, 35, 36, 37, 38, 39, 40, 41, 42, 43, 44, 45, 46, 47, 50, 51, 52, 53, 54, 55, 56, 58, 61, 63, 64, 65, 66, 67, 68, 69, 70, 71, 72, 73, 74, 76, 77, 78, 79, 80, 81, 84, 85, 86, 87, 88, 89, 90, 91, 92, 93, 94, 95, 96, 97, 98, 99, 100, 101, 102, 103, 104, 105, 106, 107, 108, 109, 110, 112, 113, 114, 115, 116, 117, 118, 119, 120, 121, 122, 123, 124, 125, 127, 128, 129, 130, 131, 132, 133, 134, 135, 136, 137, 139, 140, 141, 142, 143, 144, 145, 146, 147, 148, 150, 151, 152, 153, 154, 155, 156, 157, 158, 159, 160, 161, 162, 163, 164, 165, 166, 167, 168, 170, 171, 172, 174, 175, 176, 177, 180, 181, 182, 183, 184, 185, 186, 187, 188, 189, 190, 191, 192, 193, 194, 195, 196, 198, 200, 201, 202, 203, 205, 206, 209, 210, 211, 212, 214, 215, 217, 218, and 220.

### 2. Phenotypic Analyses

Using the original brown rust phenotypes and their normalized values for all the available individuals, we checked the residuals' distribution based on the created mixed model. With the Shapiro-Wilk test, we obtained

I. A p-value of 0.00000000000001160501 for the model built with the original values and

II. A p-value of 0.07729799 for the model built with the normalized values.

It is clear that the original dataset does not follow a normal distribution, in contrast to the normalized data. As shown in Supplementary Fig. S1, the same conclusion can be reached by visualizing the Q-Q plots. Therefore, we decided to perform our analyses using the dataset of normalized values. With these values, we estimated the BLUPs for the genotyped individuals. Using these values, we constructed a pairwise Euclidean distance matrix and performed complete hierarchical clustering to check for patterns in the dataset. There was clear separation of 2 rust phenotypic groups in the dendrogram shown in Supplementary Fig. S2.

However, to properly define these two groups, we used two different approaches: (1) the within-cluster sum of squares for different numbers of defined clusters and (2) the average silhouette width of clusters with different numbers of groups. The results are shown in Supplementary Figs. S3 and S4. For both methods, the most appropriate number of clusters was two, as expected.

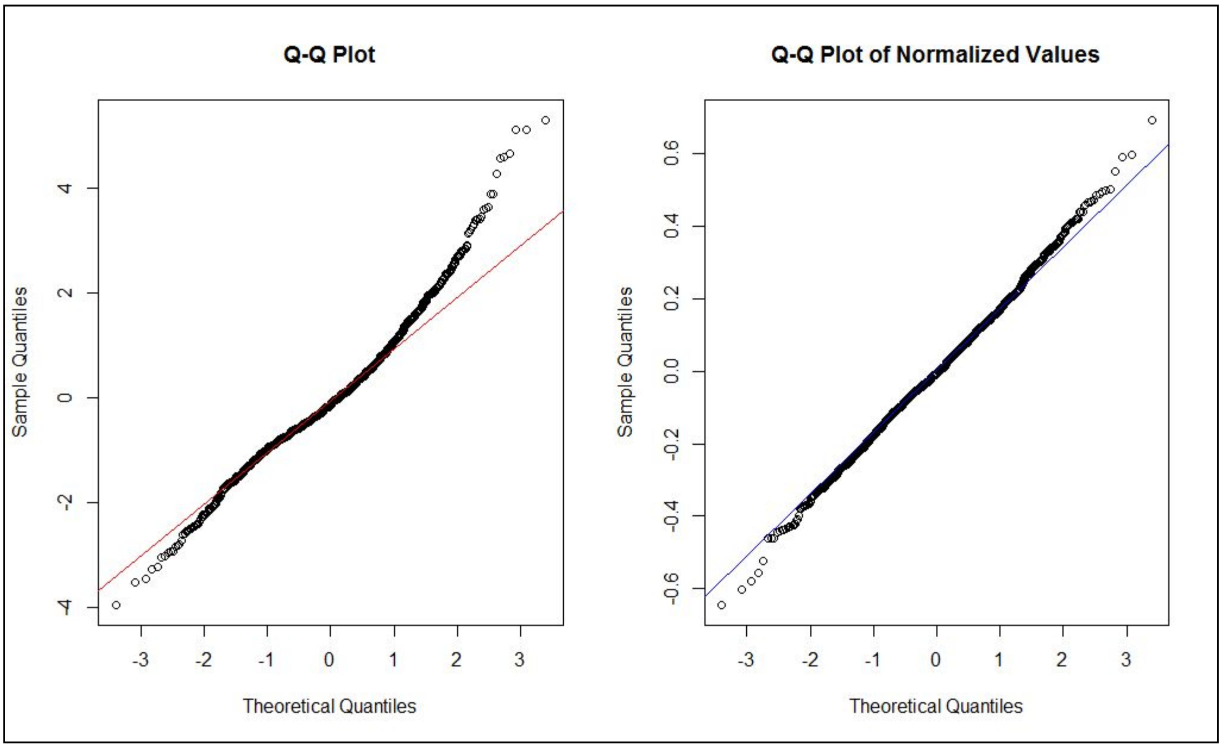


**Supplementary Fig. S1. Quantile-quantile (Q-Q) plots of brown-rust residual values.**

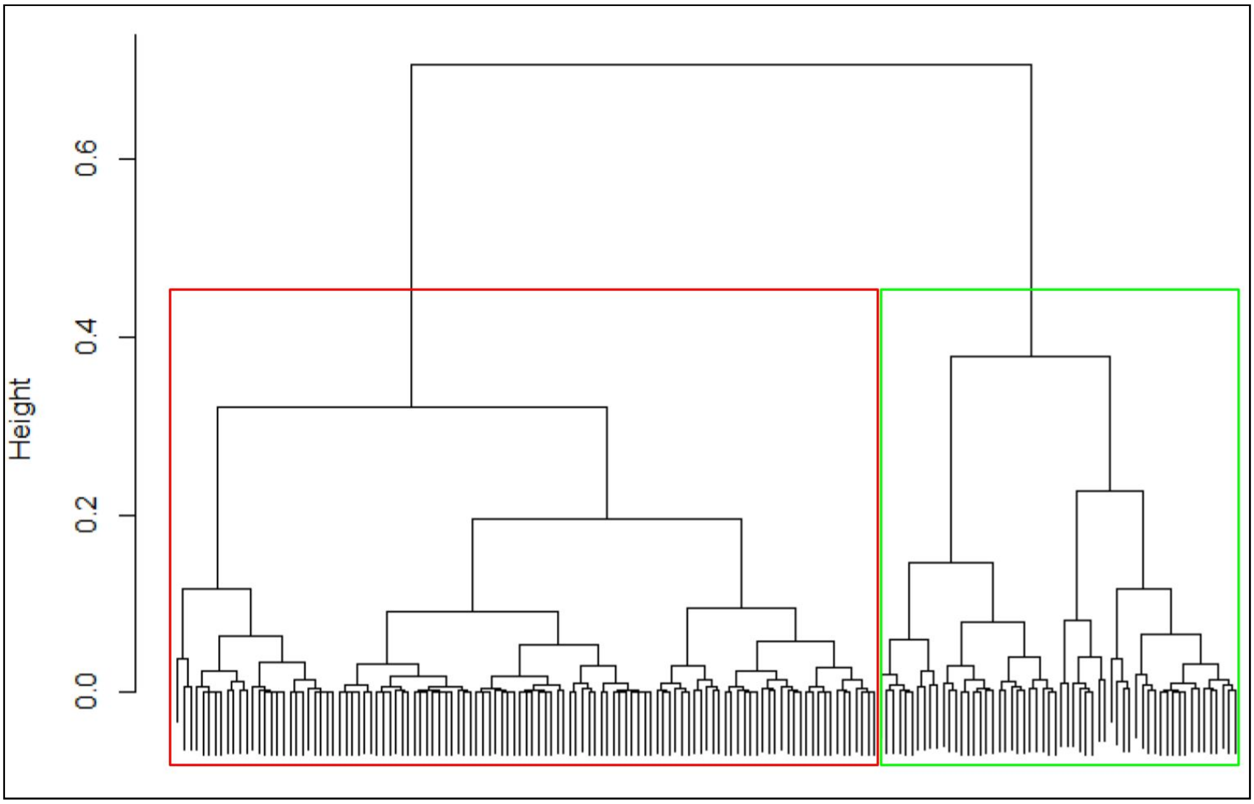


**Supplementary Fig. S2. Dendrogram based on the hierarchical clustering of Euclidean distances calculated with BLUP values obtained through a brown rust mixed model.**

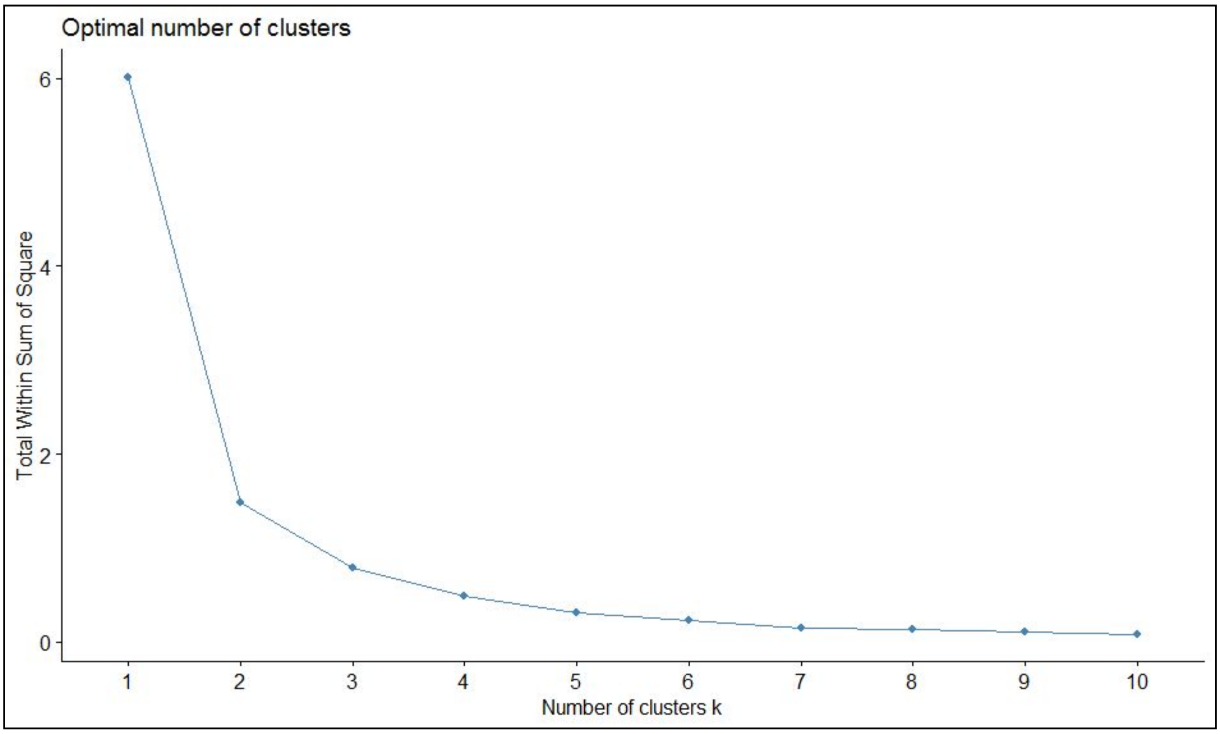


**Supplementary Fig. S3. Evaluation of the ideal quantity of clusters on the basis of within-cluster sums of squares and the BLUP values.**

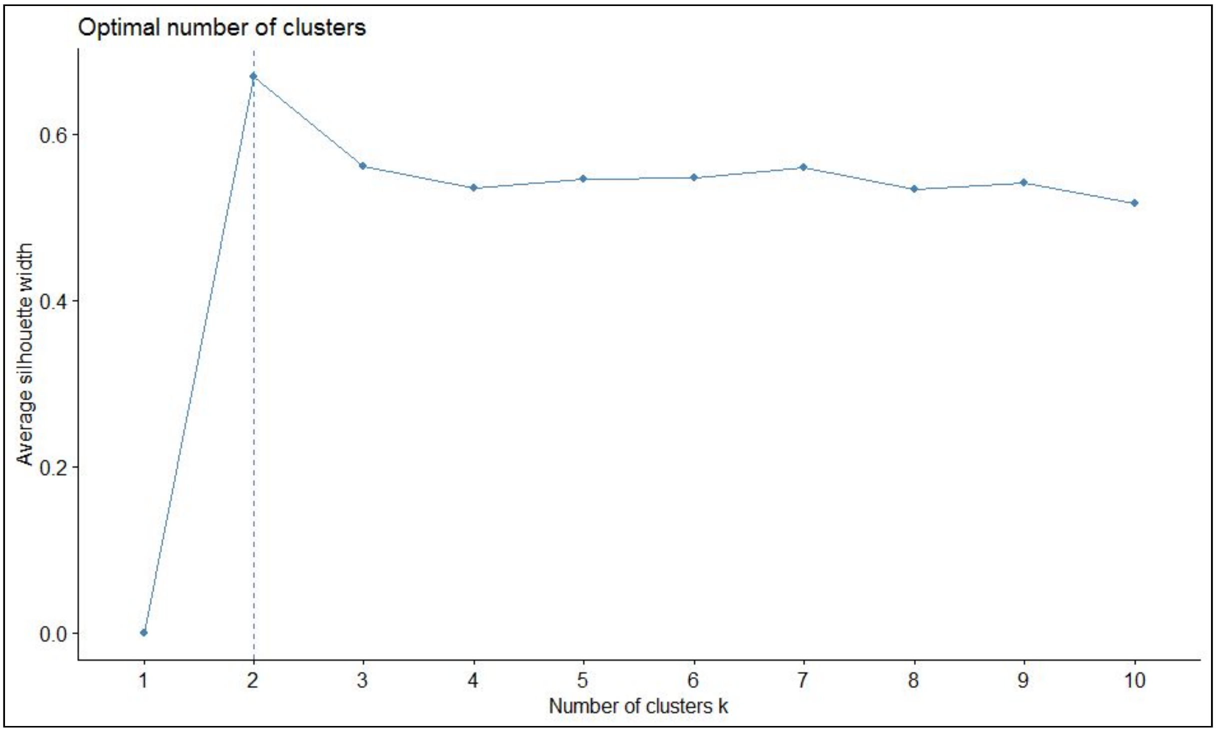


**Supplementary Fig. S4. Evaluation of the ideal quantity of clusters on the basis of average silhouette width and the BLUP values.**

The difference in BLUP values was highly variable, as shown in Supplementary Fig. S5. When evaluating all the original values without considering the BLUPs, clear differentiation was also observed (Supplementary Fig. S6), with only a few outliers. These discrepant values can be explained by the presence of nongenetic effects on the collected phenotypes. By performing a T-test between the values of these samples, we obtained a p-value of approximately 0.00000000000000022, showing significant divergence between the means.

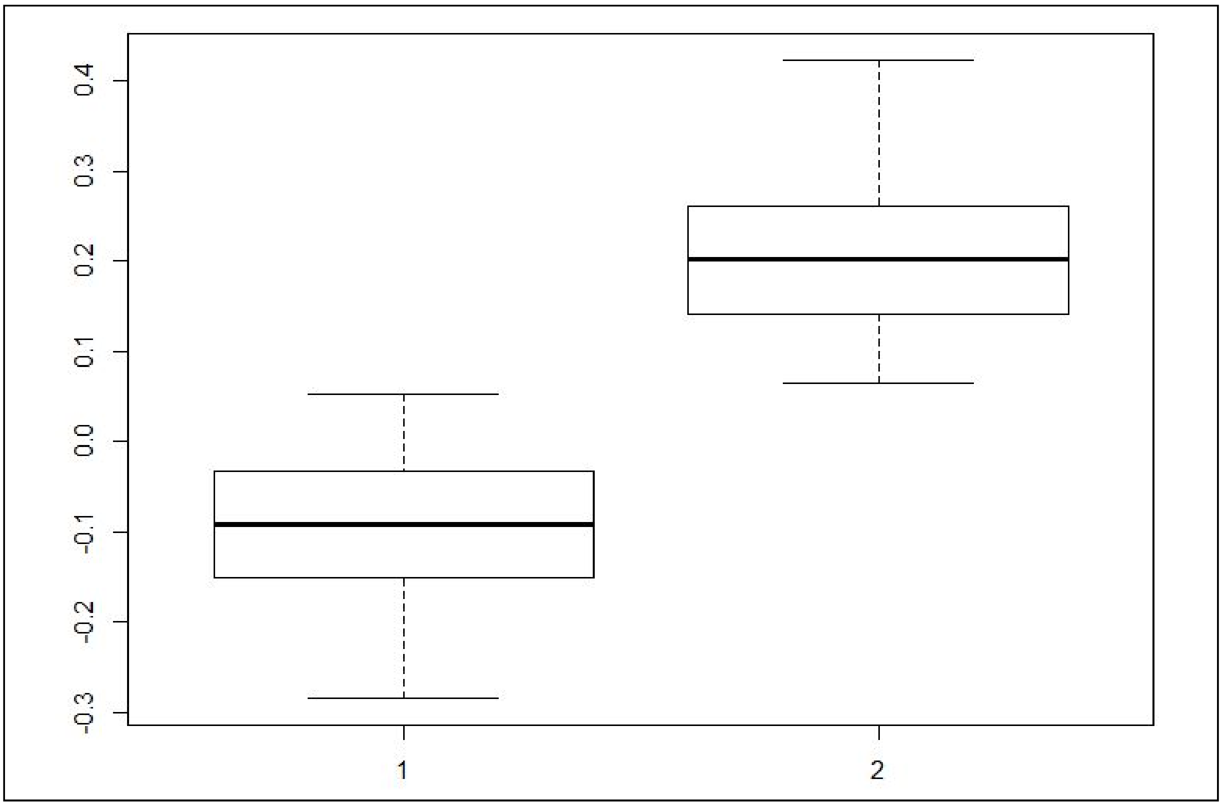


**Supplementary Fig. S5. BLUP value distributions of the identified groups.**

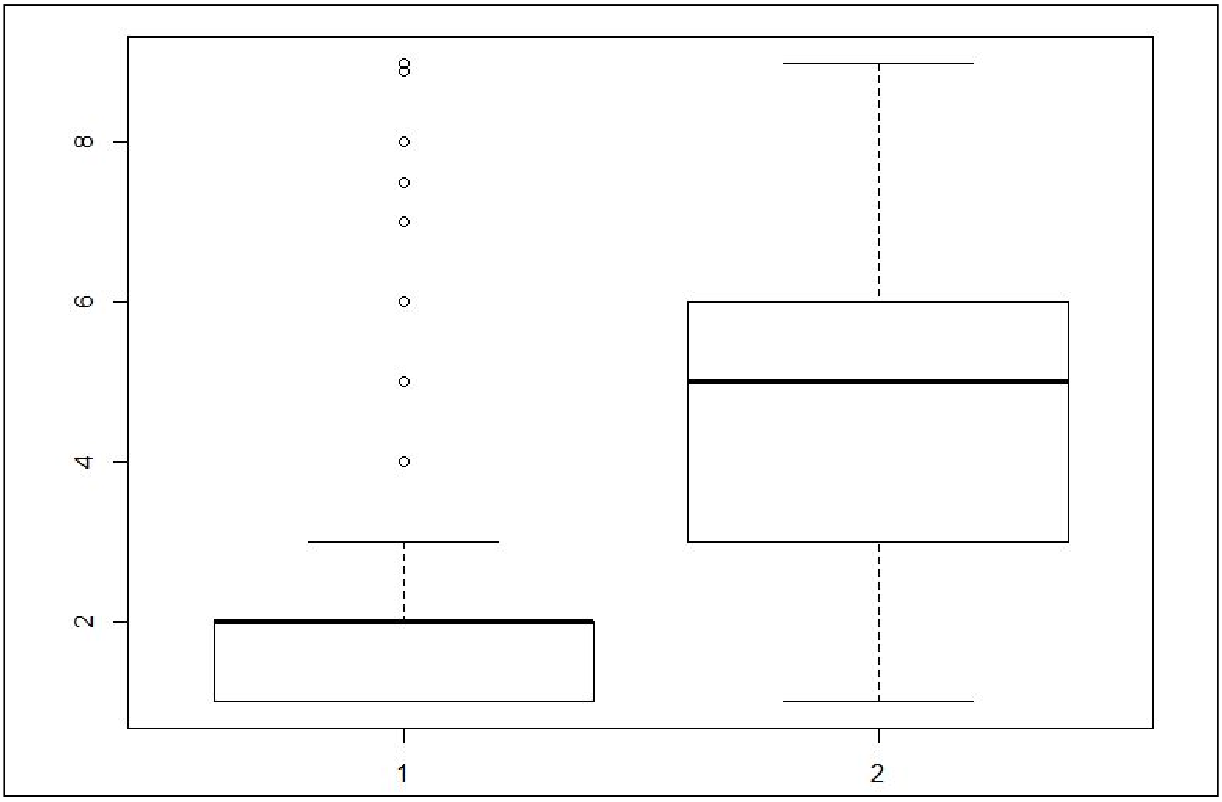


**Supplementary Fig. S6. Phenotypic value distributions of the identified groups.**

Given these findings, we continued our analyses using these rust phenotypic groups instead of the evaluation scores due to the clearly different susceptibility profiles. Therefore, we considered the individuals as belonging to group 1, named resistant, or group 2, named susceptible.

#### 3. Quality Control of Sequencing Data

Using the established quality filters on raw sequencing data, we kept 75.68% of the data (Supplementary Table S1). These data represent a subset of the original data with high-quality metrics, especially considering the demultiplexing process where there were no significant losses (Supplementary Table S2).

**Supplementary Table S1. Quantity of reads in the different GBS sequencing flow cells with and without filtering.**

| | Raw Reads | Filtered Reads |
|---|---|---|
| GBS 2015 | 164,171,329 | 134,809,753 (82.12%) |
| GBS 2017 - Exp. 1 | 149,635,834 | 85,653,323 (57.24%) |
| GBS 2017 - Exp. 2 - 1 | 341,969,762 | 278,490,414 (81.44%) |
| GBS 2017 - Exp. 2 - 2 | 311,792,539 | 257,685,281 (82.65%) |
| GBS 2017 - Exp. 3 | 135,593,786 | 78,270,511 (57.72%) |
| **Total** | **1,103,163,250** | **834,909,282 (75.68%)** |

**Supplementary Table S2. Quantity of reads kept by demultiplexing.**

| | Reads with Barcode Correspondence | Reads without Barcode Correspondence |
|---|---|---|
| GBS 2015 | 134,809,665 | 88 |
| GBS 2017 - Exp. 1 | 85,652,218 | 1,105 |
| GBS 2017 - Exp. 2 - 1 | 278,489,562 | 852 |
| GBS 2017 - Exp. 2 - 2 | 257,684,834 | 447 |
| GBS 2017 - Exp. 3 | 78,270,085 | 426 |
| **Total** | **834,906,364** | **2,918** |

#### 4. Read Alignment

The filtered read alignments against the seven selected references and the 3 adjacent subsets of the *S. spontaneum* genome obtained using BWA and Bowtie2 presented discrepancies in alignment quantity. Only uniquely mapped reads were kept for SNP calling, and the BWA results included more of these reads than did the Bowtie2 results (40.23% compared to 34.65%, respectively), as presented in Supplementary Table S3. In addition, based on the total number of aligned reads, the percentage of loss was smaller for the BWA alignments than for the Bowtie2 alignments. The mean percentage of read maintenance was approximately 10% larger with BWA than with Bowtie2. Due to this more efficient use of our data, we selected BWA alignments for SNP detection.

**Supplementary Table S3. Analysis of mapped reads using BWA and Bowtie2 against the used genomic references.**

| | **BWA Mapped Reads** | **BWA Uniquely Mapped Reads** | **Percentage of Maintenanc e** | **Bowtie Mapped Reads** | **Bowtie Uniquely Mapped Reads** | **Percentage of Maintenanc e** |
|---|---|---|---|---|---|---|
| MF genome | 787,597,773 (94.33%) | 580,694,803 (69.55%) | 73.73% | 790,307,868 (94.66%) | 415,316,587 (49.74%) | 52.55% |
| Sorghum | 372,476,439 (44.61%) | 334,543,933 (40.07%) | 89.82% | 379,731,080 (45.48%) | 312,688,984 (37.45%) | 82.34% |
| SP80-3280 | 612,219,802 (73.33%) | 256,830,971 (30.76%) | 41.95% | 626,857,213 (78.08%) | 203,710,582 (24.40%) | 32.50% |
| R570 | 453,755,780 (54.35%) | 371,364,888 (44.48%) | 81.84% | 465,348,929 (55.74%) | 339,658,623 (40.68%) | 73.00% |
| RNAseq | 286,463,784 (34.31%) | 282,308,600 (33.81%) | 98.55% | 295,867,783 (35.44%) | 279,018,734 (33.42%) | 94.31% |
| Sucest | 313,409,778 (37.54%) | 165,927,835 (19.87%) | 52.94% | 323,237,607 (38.72%) | 155,002,063 (18.57%) | 47.95% |
| SS A1 | 595,267,463 (71.30%) | 342,151,259 (41.00%) | 57.48% | 609,900,489 (73.05%) | 298,041,048 (35.70%) | 48.87% |
| SS A2 | 594,220,509 (71.17%) | 346,818,994 (41.54%) | 58.37% | 609,010,287 (72.94%) | 301,122,972 (36.07%) | 49.44% |
| SS A3 | 576,398,435 (69.04%) | 336,411,572 (40.30%) | 58.37% | 590,092,154 (70.68%) | 294,076,883 (35.22%) | 49.84% |
| SS A4 | 587,266,198 (70.34%) | 341,480,330 (40.90%) | 58.15% | 599,587,919 (71.81%) | 293,878,819 (35.20%) | 49.01% |
| **Mean** | **62.03%** | **40.23%** | **67.12%** | **63.66%** | **34.65%** | **57.98%** |

Supplementary Fig. S7 provides an overview of the alignment process across the different references. Due to very low sequencing depths, individuals 33, 39, 41, 47, 51, 56, 68, 85, 87, 93 and 107 were not considered in further analyses (Supplementary Fig. S7-b). Either on different scales (original and logarithmic) or with different references, the discrepancy in these observations when compared to the entire dataset was evident, and bias in further analyses could potentially be introduced if using these individuals, such as filtering SNP positions, due to the small number of individuals. The remaining individuals were classified into rust phenotypic groups, with 118 individuals classified as belonging to the "resistant" group and 53 as belonging to the "susceptible" group.

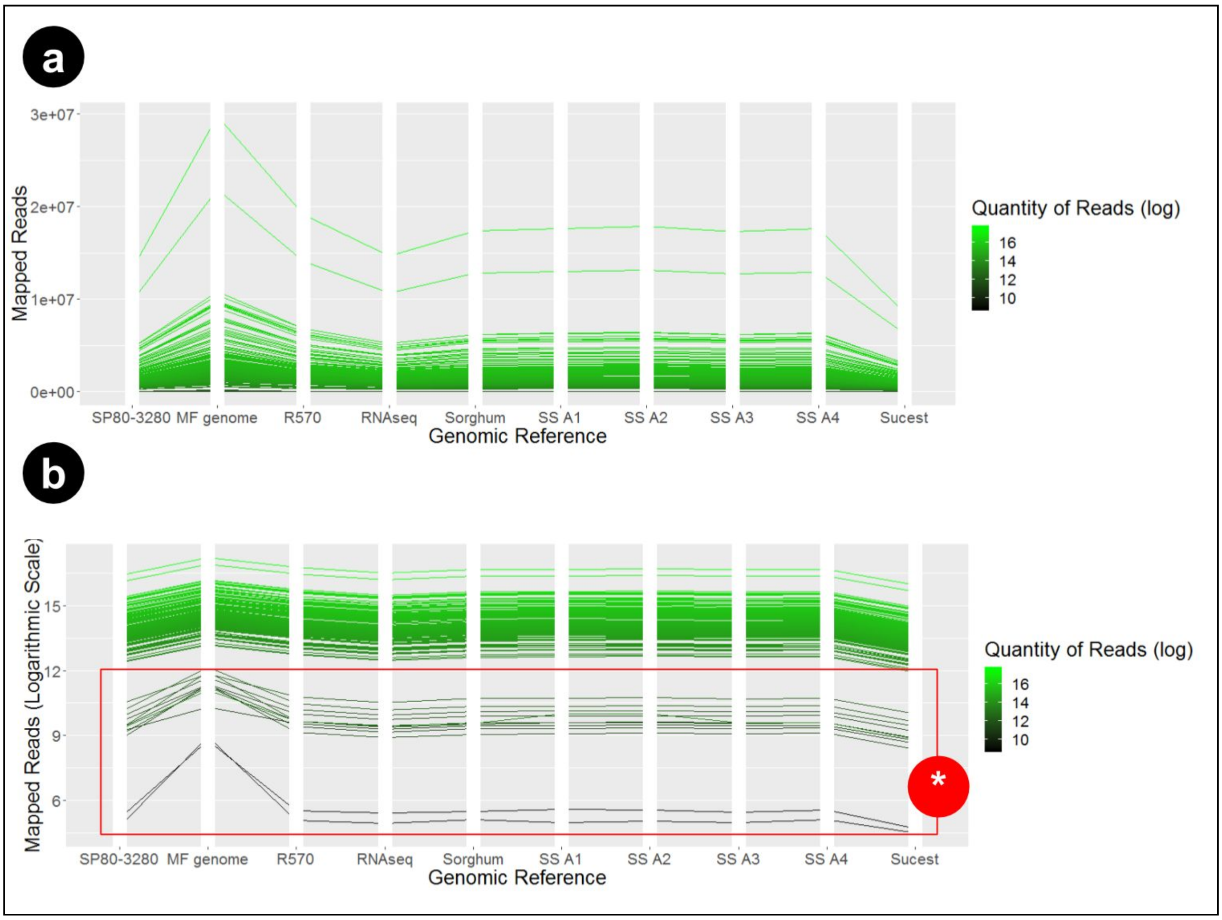


**Supplementary Fig. S7. (a) Quantity of mapped reads per individual; (b) quantity of mapped reads per individual on a logarithmic scale; (*) individuals removed from the analyses.**

#### 5. Reference Evaluation

To evaluate the reads' correspondence to the different references used, we compared the results shown in Supplementary Tables S4 and S5. Although we had references containing more information than the MF genome contained (Supplementary Table S4), we obtained more extensive alignments of our data to the MF reference (Supplementary Table S5), which was built according to the obtained consensus contigs. The total length of MF contigs was at least 6.5 times greater than that for the other references, and the number of contigs was at least 6.6 times higher, indicating a larger percentage of genomic regions with similarity to the GBS data. Although the N50 metric for the original MF scaffolds was lower than that for the other references, the value had another interpretation when using the consensus sequences, where all the regions had similar values. However, analysis of the L50 metric together with the total quantity of contigs corroborated that the MF alignments produced longer sequences.

**Supplementary Table S4. Analysis of different reference compositions.**

| | Total (bp) | Ns per 100 kbp | No. Contigs or Scaffolds | Largest Contig (bp) | N50 (bp) |
|---|---|---|---|---|---|
| MF genome | 674,032,540 | 46,984.06 | 1,109,444 | 35,917 | 2,083 |
| Sorghum | 708,863,705 | 4,720.34 | 870 | 80,884,392 | 68,658,214 |
| SP80-3280 | 1,169,948,913 | 0.00 | 199,028 | 115,913 | 8,451 |
| R570 | 426,798,785 | 8,814.84 | 211 | 72,286,729 | 45,576,573 |
| RNAseq | 66,572,642 | 0.00 | 72,269 | 17,055 | 1,737 |
| Sucest | 133,946,191 | 0.00 | 195,765 | 8,854 | 1,258 |
| SS A1 | 974,684,959 | 198.19 | 15,744 | 121,903,222 | 79,322,609 |
| SS A2 | 984,574,931 | 198.97 | 15,744 | 123,268,843 | 90,128,929 |
| SS A3 | 963,861,475 | 197.93 | 15,744 | 126,636,275 | 89,762,077 |
| SS A4 | 938,617,136 | 194.65 | 15,744 | 116,847,860 | 84,617,075 |

**Supplementary Table S5. Consensus sequence profiles obtained via read mapping against the different references.**

| | Contigs | Total (bp) | Ns per 100 kbp | Largest Contig (bp) | N50/N75 | L50/L75 |
|---|---|---|---|---|---|---|
| MF genome | 82,472 | 7,087,591 | 60.51 | 120 | 86/86 | 40,353/ 60,957 |
| Sorghum | 35,174 | 3,025,392 | 474.38 | 123 | 86/86 | 17,058/ 25,852 |
| SP80-3280 | 43,639 | 3,759,046 | 80.66 | 123 | 86/86 | 21,414/ 32,342 |
| R570 | 48,663 | 4,203,728 | 92.13 | 118 | 86/86 | 23,829/ 36,049 |
| RNAseq | 25,673 | 2,205,254 | 73.01 | 116 | 86/86 | 12,536/ 18,946 |
| Sucest | 15,335 | 1,311,672 | 77.76 | 120 | 86/86 | 7,466/ 11,279 |
| SS A1 | 49,487 | 4,271,185 | 131.44 | 124 | 86/86 | 24,185/ 36,602 |
| SS A2 | 49,616 | 4,281,974 | 126.95 | 121 | 86/86 | 24,239/ 36,687 |
| SS A3 | 47,994 | 4,142,664 | 130.23 | 122 | 86/86 | 23,453/ 35,496 |
| SS A4 | 48,600 | 4,195,855 | 138.90 | 121 | 86/86 | 23,744/ 35,941 |

It might be intuitive to conclude that with more extensive alignments (i.e., greater coverage), the sequencing depths across MF scaffolds would be lower.

However, this was not observed. MF alignments produced a larger number of genomic positions with high correspondence than did other alignments (Supplementary Table S5). However, as shown in Supplementary Table S6, these positions also had increased depth. With at least 44.38% more coverage than the other references, MF alignments had a larger quantity of reads corroborating genomic positions. Considering a minimum depth of 8.000 reads, MF had at least 64.50% more covered regions than the other references.

**Supplementary Table S6. Depth mapping profiles of different references and GBS data.**

| | Total Positions (bp) | Total Positions where Mapping Depth > Value | | | | | |
|---|---|---|---|---|---|---|---|
| | | >=100 | >=500 | >=1000 | >=5000 | >=6000 | >=8000 |
| MF genome | 13,694,206 | 5,630,873 | 3,855,721 | 3,060,666 | 1,343,310 | 1,172,174 | 739,799 |
| Sorghum | 6,577,944 | 2,750,262 | 1,961,559 | 1,594,831 | 773,279 | 683,861 | 438,040 |
| SP80-3280 | 8,119,350 | 2,881,841 | 1,915,561 | 1,480,226 | 615,468 | 536,882 | 336,060 |
| R570 | 8,300,203 | 3,236,343 | 2,234,123 | 1,788,540 | 822,918 | 722,547 | 449,732 |
| RNAseq | 4,482,524 | 1,978,380 | 1,414,520 | 1,150,812 | 558,839 | 494,594 | 317,338 |
| Sucest | 2,857,978 | 1,248,375 | 886,462 | 719,308 | 342,231 | 305,128 | 199,139 |
| SS A1 | 9,464,530 | 3,339,323 | 2,256,387 | 1,778,348 | 789,632 | 690,830 | 427,588 |
| SS A2 | 9,485,051 | 3,358,500 | 2,282,746 | 1,810,360 | 812,133 | 708,668 | 436,856 |
| SS A3 | 9,254,628 | 3,243,610 | 2,197,171 | 1,740,758 | 776,469 | 680,257 | 421,163 |
| SS A4 | 9,282,252 | 3,273,861 | 2,221,450 | 1,754,522 | 782,224 | 685,346 | 425,154 |

The MF sugarcane genome had the largest number of total mapped positions. Additionally, this reference also had the largest number of mapped positions in five of the six evaluated depth ranges. Importantly, in addition to having the greatest observed depths, these reads were better distributed among the scaffolds, as shown by the quantity of mapped positions. On the basis of these conclusions, we evaluated redundancy among the consensus sequences obtained by the mapping process against the different references. Supplementary Table S7 and Supplementary Fig. S8 show this redundancy. MF contigs had the largest values of correspondence, i.e., they included the majority of genomic information contained in all other references.

**Supplementary Table S7. Consensus sequence redundancies across different references assessed via comparative alignments (alignment queries as rows and targets as columns).**

| | MF genome | Sorghum | SP80-3280 | R570 | RNAseq | Sucest | SS A1 | SS A2 | SS A3 | SS A4 |
|---|---|---|---|---|---|---|---|---|---|---|

| | | | | | | | | | | |
|---|---|---|---|---|---|---|---|---|---|---|
| MF genome | 82,472 (100%) | 25,072 (71.28%) | 27,733 (78.37%) | 35,249 (72.43%) | 20,093 (78.37%) | 11,670 (76.10%) | 33,509 (67.71%) | 33,925 (68.38%) | 32,650 (68.03%) | 33,087 (68.08%) |
| Sorghum | 24,647 (29.89%) | 35,174 (100%) | 10,124 (23.20%) | 16,963 (34.86%) | 11,927 (46.46%) | 6,613 (43.13%) | 14,792 (29.90%) | 15,087 (30.41%) | 14,634 (30.49%) | 14,700 (30.25%) |
| SP80-3280 | 27,825 (33.74%) | 10,340 (29.40%) | 43,639 (100%) | 16,418 (33.74%) | 8,935 (34.80%) | 5,216 (34.01%) | 16,474 (33.29%) | 16,556 (33.37%) | 15,792 (32.90%) | 15,989 (32.90%) |
| R570 | 35,078 (42.53%) | 17,163 (48.80%) | 16,206 (37.14%) | 48,663 (100%) | 12,650 (49.27%) | 7,247 (47.26%) | 21,321 (43.08%) | 21,362 (43.05%) | 21,135 (44.04%) | 21,319 (43.87%) |
| RNAseq | 19,634 (23.81%) | 11,851 (33.69%) | 8,703 (19.94%) | 12,426 (25.53%) | 25,673 (100%) | 8,328 (54.31%) | 11,115 (22.46%) | 11,199 (22.57%) | 10,902 (22.72%) | 11,148 (22.94%) |
| Sucest | 11,380 (13.80%) | 6,590 (18.74%) | 5,068 (11.61%) | 7,115 (14.62%) | 8,325 (32.43%) | 15,335 (100%) | 6,623 (13.38%) | 6,601 (13.30%) | 6,324 (13.18%) | 6,542 (13.46%) |
| SS A1 | 33,196 (40.25%) | 14,940 (42.47%) | 16,196 (37.11%) | 21,255 (43.68%) | 11,260 (43.86%) | 6,729 (43.88%) | 49,487 (100%) | 25,256 (50.90%) | 24,370 (50.78%) | 24,837 (51.10%) |
| SS A2 | 33,596 (40.74%) | 15,199 (43.21%) | 16,235 (37.20%) | 21,306 (43.78%) | 11,368 (44.28%) | 6,690 (43.63%) | 25,227 (50.98%) | 49,616 (100%) | 24,599 (51.25%) | 24,905 (51.24%) |
| SS A3 | 32,321 (39.19%) | 14,731 (41.88%) | 15,526 (35.58%) | 21,092 (43.34%) | 11,049 (43.04%) | 6,410 (41.80%) | 24,330 (49.16%) | 24,612 (49.60%) | 47,994 (100%) | 23,789 (48.95%) |
| SS A4 | 32,797 (39.77%) | 14,837 (42.18%) | 15,697 (35.97%) | 21,286 (43.74%) | 11,316 (44.08%) | 6,654 (43.39%) | 24,866 (50.25%) | 24,918 (50.22%) | 23,855 (49.70%) | 48,600 (100%) |

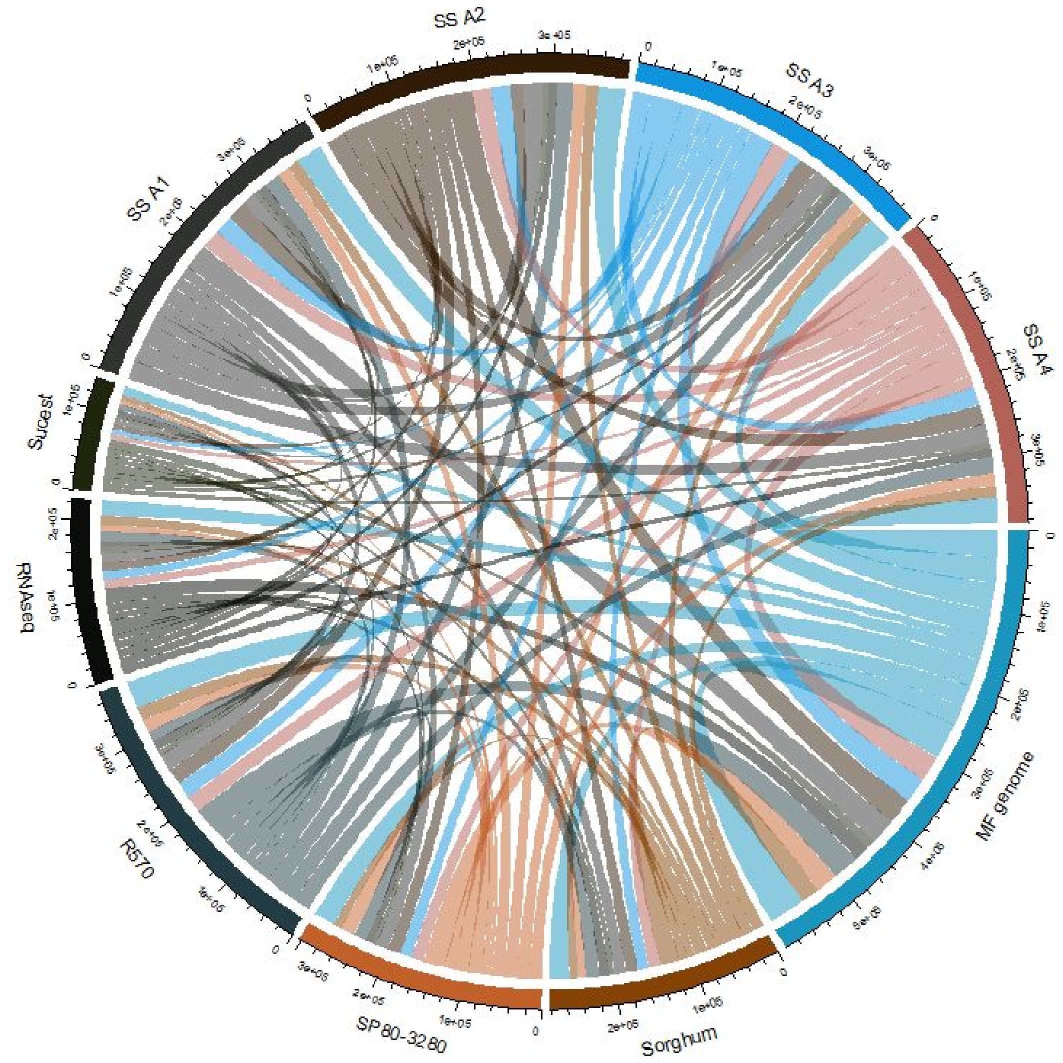


**Supplementary Fig. S8. Redundancies of consensus sequences visualized in a Circos plot.**

As a final step in considering the MF genome as our most appropriate option for the analyses, we evaluated the quantity of SNPs identified by Tassel and Stacks, tools developed to identify these variants in GBS and RADSeq data. Due to the specificity of these experiments in selecting genomic regions, these tools were selected as another reference evaluation parameter. As shown in Supplementary Table S8, raw variants called by Tassel and the MF reference genome allowed the identification of at least 51% more variants than did those obtained with the other references.

**Supplementary Table S8. Different variants identified by Tassel.**

| | Raw Variants | Biallelic SNPs | Multiallelic SNPs | Indels |
|---|---|---|---|---|
| MF genome | 137,757 | 135,594 | 385 | 1,778 |
| Sorghum | 63,653 | 62,467 | 105 | 1,081 |
| SP80-3280 | 63,625 | 62,781 | 208 | 636 |
| R570 | 91,108 | 89,795 | 147 | 1,166 |
| RNAseq | 60,214 | 59,291 | 57 | 866 |
| Sucest | 38,036 | 37,363 | 70 | 603 |
| SS A1 | 86,634 | 85,419 | 171 | 1,044 |
| SS A2 | 89,038 | 87,728 | 225 | 1,085 |
| SS A3 | 85,306 | 84,119 | 157 | 1,030 |
| SS A4 | 86,563 | 85,358 | 176 | 1,029 |

The same result was observed when using Stacks (Supplementary Table S9). The number of identified variants was much larger when using the MF reference than when using the other references, even when using the established filters.

**Supplementary Table S9. Identification of SNPs using Stacks and Tassel and stringent filters (at least 50 reads per individual and a maximum of 25% missing data)**

| | # of Tassel Biallelic SNPs | # of Tassel Filtered Biallelic SNPs | # of Stacks Biallelic SNPs | # of Stacks Filtered Biallelic SNPs | # Intersecting Biallelic SNPs |
|---|---|---|---|---|---|
| MF genome | 137,757 | 18,881 | 106,881 | 2,605 | 50,274 |
| Sorghum | 63,653 | 10,262 | 44,074 | 1,544 | 16,159 |
| SP80-3280 | 63,624 | 7,552 | 56,830 | 1,033 | 25,162 |
| R570 | 91,108 | 13,441 | 72,378 | 1,784 | 33,752 |
| RNAseq | 60,214 | 10,881 | 38,641 | 1,296 | 18,715 |
| Sucest | 38,036 | 6,422 | 23,545 | 623 | 11,001 |

| | | | | | |
|---|---|---|---|---|---|
| SS A1 | 86,634 | 11,536 | 72,638 | 1,518 | 32,570 |
| SS A2 | 89,038 | 11,763 | 73,536 | 1,600 | 33,098 |
| SS A3 | 85,306 | 11,726 | 70,938 | 1,492 | 31,612 |
| SS A4 | 86,563 | 11,562 | 71,539 | 1,502 | 31,876 |

### 6. SNP Calling

SNP calling was also performed with GATK, FreeBayes and SAMtools. The tools produced different quantities of identified markers, confirming the influence of the SNP caller on the inference of putative variants. Due to sugarcane's aneuploidy at the locus level, it was not possible to define an exclusive ploidy to be used for the entire dataset. Due to this difficulty, we tested all the possible ploidy configurations with GATK and FreeBayes to assess the influence of this parameter on the final set. The quantities of markers were similar (Supplementary Table S10). Furthermore, these positions were the same for different ploidies, with only the quantity of detected variants and related allelic dosages changing.

**Supplementary Table S10. Identification of SNPs using the MF genome, GATK, FreeBayes and SAMtools.**

| | GATK | FreeBayes | SAMtools |
|---|---|---|---|
| Ploidy 2 | 61,604 | 77,079 | 353,715 |
| Ploidy 4 | 61,481 | 77,079 | - |
| Ploidy 6 | 61,441 | 77,079 | - |
| Ploidy 8 | 61,317 | 77,079 | - |
| Ploidy 10 | 62,083 | 76,932 | - |
| Ploidy 12 | 62,059 | 76,932 | - |
| Ploidy 14 | 0 | 76,560 | - |
| Ploidy 16 | 0 | 76,008 | - |
| Ploidy 18 | 0 | 73,110 | - |
| Ploidy 20 | 0 | 72,391 | - |

Although there was only a small discrepancy in the positions of the analyzed variants among ploidy configurations, the quantities of missing data were very different (Supplementary Table S11). As the defined ploidy level increased, the quantity of missing data also increased. This fact may be explained by the depth required by these tools to characterize a haplotype. To increase the reliability of the dataset, the intersection of these positions between different ploidy configurations (2-12) was examined, and individual genotypes were further evaluated.

**Supplementary Table S11. Different missing data distributions across the ploidy range used.**

| Ploidy | Tool | SNPs | Maximum of 90% NAs | Maximum of 75% NAs | Maximum of 50% NAs | Maximum of 25% NAs |
|---|---|---|---|---|---|---|
| 2 | GATK | 61,604 | 40,486 | 35,947 | 27,504 | 16,578 |
| | FreeBayes | 77,079 | 73,668 | 69,650 | 60,335 | 49,492 |
| | SAMtools | 353,715 | 227,844 | 195,180 | 156,848 | 123,678 |
| 4 | GATK | 61,481 | 40,441 | 35,901 | 27,463 | 16,548 |
| | FreeBayes | 77,079 | 73,668 | 69,650 | 60,335 | 49,492 |
| 6 | GATK | 61,441 | 40,413 | 35,865 | 27,427 | 16,531 |
| | FreeBayes | 77,079 | 73,668 | 69,650 | 60,335 | 49,492 |
| 8 | GATK | 61,317 | 40,347 | 35,803 | 27,373 | 16,491 |
| | FreeBayes | 77,079 | 73,668 | 69,650 | 60,335 | 49,492 |
| 10 | GATK | 62,083 | 25,861 | 16,251 | 11,190 | 7,543 |
| | FreeBayes | 76,932 | 73,527 | 69,526 | 60,236 | 49,412 |
| 12 | GATK | 62,059 | 25,882 | 16,260 | 11,200 | 7,546 |
| | FreeBayes | 76,932 | 73,527 | 69,526 | 60,236 | 49,412 |

### 7. Cross-Validation Results with Allelic Dosages

The selection of markers based on the defined criteria for predicting rust phenotypic groups was organized into allelic dosages (ADs) and allelic proportions (APs) with the following subsets:

1. All (total SNP dataset);
2. FS1 (feature selection 1 - gradient tree boosting);
3. FS2 (feature selection 2 - L1-based SVC);
4. FS3 (feature selection 3 - extra trees);
5. FS4 (feature selection 4 - F statistic from ANOVA); and
6. FS5 (feature selection 5 - random forests).

Accuracy, recall, precision and specificity metrics for each subset of markers are provided in Supplementary Tables S12 to S17. The first comparison was performed to evaluate the appropriateness of using the ADs or APs for this task. For each measure, we calculated the percentage increase caused by using APs instead of ADs, coloring these percentages blue (increase), red (decrease) or black (maintenance).

**Supplementary Table S12. Cross-validation results when using the entire dataset.**

| Performances | | | | | | | | |
|---|---|---|---|---|---|---|---|---|
| Model | Accuracy (AD) | Accuracy (AP) | Recall (AD) | Recall (AP) | Precision (AD) | Precision (AP) | Specificity (AD) | Specificity (AP) |
| AB | 52.72 | 58.75 **(+11%)** | 53.69 | 60.14 **(+12%)** | 70.74 | 75.12 **(+6%)** | 50.55 | 55.66 **(+10%)** |
| DT | 51.95 | 55.08 **(+6%)** | 53.18 | 55.32 **(+4%)** | 69.99 | 73.04 **(+4%)** | 49.23 | 54.53 **(+11%)** |
| GP | 31.27 | 68.94 **(+20%)** | 0.41 | 99.80 **(+24241%)** | 100.00 | 69.01 **(-31%)** | 100.00 | 0.23 **(-100%)** |
| KNN | 68.97 | 68.60 **(-1%)** | 99.95 | 98.97 **(-1%)** | 68.99 | 69.00 **(+0%)** | 0.00 | 1.00 **(+0%)** |
| MLP | 54.58 | 49.26 **(-10%)** | 59.69 | 48.14 **(-19%)** | 70.05 | 68.96 **(-2%)** | 43.19 | 51.75 **(+20%)** |
| GNB | 50.33 | 47.85 **(-5%)** | 42.36 | 39.47 **(-7%)** | 74.70 | 72.40 **(-3%)** | 68.06 | 66.49 **(-2%)** |
| RF | 48.55 | 50.12 **(+3%)** | 41.64 | 43.70 **(+4%)** | 71.99 | 73.21 **(+2%)** | 63.92 | 64.40 **(+0%)** |
| SVM | 47.61 | 53.91 **(+13%)** | 35.99 | 52.87 **(+47%)** | 75.13 | 72.89 **(-3%)** | 73.47 | 56.21 **(-23%)** |
| Mean | 51.14 | 54.50 **(+7%)** | 47.77 | 54.10 **(+13%)** | 71.37 | 72.65 **(+2%)** | 57.24 | 55.10 **(-4%)** |

**Supplementary Table S13. Cross-validation results when using the dataset obtained with gradient tree boosting.**

| Performances | | | | | | | | |
|---|---|---|---|---|---|---|---|---|
| Model | Accuracy (AD) | Accuracy (AP) | Recall (AD) | Recall (AP) | Precision (AD) | Precision (AP) | Specificity (AD) | Specificity (AP) |
| AB | 71.49 | 77.50 **(+8%)** | 72.36 | 78.07 **(+8%)** | 84.09 | 87.97 **(+4%)** | 69.53 | 76.23 **(+10%)** |
| DT | 61.20 | 66.23 **(+8%)** | 61.56 | 67.12 **(+9%)** | 77.58 | 80.70 **(+4%)** | 60.40 | 64.26 **(+6%)** |
| GP | 69.83 | 81.11 **(+16%)** | 98.49 | 88.63 **(-10%)** | 70.00 | 84.71 **(+21%)** | 6.02 | 64.38 **(+969%)** |
| KNN | 69.24 | 70.27 **(+1%)** | 99.78 | 98.00 **(-2%)** | 69.23 | 70.46 **(+2%)** | 1.25 | 8.53 **(+582%)** |
| MLP | 83.13 | 78.26 **(-6%)** | 86.13 | 80.49 **(-7%)** | 89.07 | 87.03 **(-2%)** | 76.47 | 73.30 **(-4%)** |
| GNB | 83.83 | 86.37 **(+3%)** | 79.87 | 83.60 **(+5%)** | 96.03 | 96.15 **(+0%)** | 92.64 | 92.55 **(+0%)** |
| RF | 63.29 | 68.60 **(+8%)** | 57.38 | 63.42 **(+11%)** | 84.43 | 87.66 **(+4%)** | 76.43 | 80.13 **(+5%)** |

| | | | | | | | | |
|---|---|---|---|---|---|---|---|---|
| SVM | 70.56 | 78.11 **(+10%)** | 61.19 | 80.59 **(+32%)** | 94.08 | 86.75 **(-8%)** | 91.43 | 72.58 **(-21%)** |
| Mean | 70.20 | 77.80 **(+11%)** | 76.12 | 80.54 **(+6%)** | 84.26 | 86.89 **(+3%)** | 72.98 | 72.94 **(+0%)** |

**Supplementary Table S14. Cross-validation results when using the dataset obtained with L1-based SVC.**

| **Performances** | | | | | | | | |
|---|---|---|---|---|---|---|---|---|
| Model | Accuracy (AD) | Accuracy (AP) | Recall (AD) | Recall (AP) | Precision (AD) | Precision (AP) | Specificity (AD) | Specificity (AP) |
| AB | 70.85 | 75.98 **(+7%)** | 71.07 | 76.59 **(+8%)** | 84.22 | 87.04 **(+3%)** | 70.36 | 74.60 **(+6%)** |
| DT | 57.18 | 60.69 **(+6%)** | 57.45 | 61.05 **(+6%)** | 74.66 | 77.21 **(+3%)** | 56.58 | 59.89 **(+6%)** |
| GP | 69.53 | 88.03 **(+27%)** | 96.61 | 99.80 **(+3%)** | 70.32 | 85.34 **(+21%)** | 9.23 | 61.83 **(+570%)** |
| KNN | 71.29 | 71.11 **(+0%)** | 98.79 | 99.22 **(+0%)** | 70.98 | 70.72 **(+0%)** | 10.06 | 8.53 **(-20%)** |
| MLP | 90.42 | 97.71 **(+8%)** | 91.56 | 98.55 **(+8%)** | 94.39 | 98.14 **(+4%)** | 87.89 | 95.85 **(+9%)** |
| GNB | 86.26 | 88.53 **(+3%)** | 86.79 | 87.15 **(+0%)** | 92.83 | 95.84 **(+3%)** | 85.08 | 91.58 **(+8%)** |
| RF | 60.75 | 66.02 **(+9%)** | 54.12 | 60.29 **(+11%)** | 83.12 | 86.35 **(+4%)** | 75.53 | 78.77 **(+4%)** |
| SVM | 49.95 | 92.50 **(+85%)** | 29.95 | 92.05 **(+207%)** | 92.37 | 96.93 **(+5%)** | 94.49 | 93.50 **(-1%)** |
| Mean | 70.19 | 82.01 **(+17%)** | 78.93 | 89.60 **(+14%)** | 83.67 | 86.70 **(+4%)** | 72.95 | 76.69 **(+5%)** |

**Supplementary Table S15. Cross-validation results when using the dataset obtained with extra trees.**

| **Performances** | | | | | | | | |
|---|---|---|---|---|---|---|---|---|
| Model | Accuracy (AD) | Accuracy (AP) | Recall (AD) | Recall (AP) | Precision (AD) | Precision (AP) | Specificity (AD) | Specificity (AP) |
| AB | 58.32 | 62.71 **(+8%)** | 58.97 | 65.08 **(+10%)** | 75.27 | 77.29 **(+3%)** | 56.87 | 57.43 **(+1%)** |
| DT | 53.65 | 56.06 **(+4%)** | 54.29 | 57.12 **(+5%)** | 71.68 | 73.31 **(+2%)** | 52.25 | 53.70 **(+3%)** |
| GP | 36.86 | 69.01 **(+87%)** | 8.97 | 100.00 **(+1015%)** | 95.06 | 69.01 **(-27%)** | 98.96 | 0.00 **(-100%)** |
| KNN | 68.96 | 69.01 **(+0%)** | 99.91 | 99.77 **(+0%)** | 69.00 | 69.07 **(+0%)** | 0.06 | 0.53 **(+783%)** |
| MLP | 73.01 | 52.99 **(-27%)** | 82.44 | 52.31 **(-37%)** | 79.28 | 71.90 **(-9%)** | 52.02 | 54.49 **(+5%)** |

| | | | | | | | | |
|---|---|---|---|---|---|---|---|---|
| GNB | 67.16 | 65.51 (-2%) | 60.49 | 58.06 (-4%) | 88.21 | 87.83 (+0%) | 82.00 | 82.09 (+0%) |
| RF | 52.53 | 55.36 (+5%) | 45.70 | 48.81 (+7%) | 75.93 | 78.34 (+3%) | 67.74 | 69.94 (+3%) |
| SVM | 61.05 | 68.27 (+12%) | 51.76 | 68.08 (+32%) | 86.32 | 82.88 (-4%) | 81.74 | 68.70 (-16%) |
| Mean | 59.69 | 64.11 (+7%) | 56.63 | 61.57 (+9%) | 77.61 | 75.3 (-3%) | 62.31 | 55.96 (-10%) |

**Supplementary Table S16. Cross-validation results when using the dataset obtained with the F statistic from ANOVA.**

| Performances | | | | | | | | |
|---|---|---|---|---|---|---|---|---|
| Model | Accuracy (AD) | Accuracy (AP) | Recall (AD) | Recall (AP) | Precision (AD) | Precision (AP) | Specificity (AD) | Specificity (AP) |
| AB | 72.37 | 73.57 (+2%) | 72.90 | 74.86 (+3%) | 84.92 | 85.05 (+0%) | 71.19 | 70.70 (-1%) |
| DT | 58.54 | 60.27 (+3%) | 58.69 | 60.85 (+4%) | 75.77 | 76.77 (+1%) | 58.21 | 59.00 (+1%) |
| GP | 69.68 | 81.64 (+17%) | 97.53 | 99.04 (+2%) | 70.16 | 79.43 (+13%) | 7.66 | 42.91 (+460%) |
| KNN | 72.18 | 72.53 (+0%) | 99.25 | 99.46 (+0%) | 71.50 | 71.70 (+0%) | 11.91 | 12.58 (+6%) |
| MLP | 95.35 | 86.81 (-9%) | 96.13 | 87.91 (-9%) | 97.10 | 92.60 (-5%) | 93.60 | 84.36 (-10%) |
| GNB | 92.89 | 90.77 (-2%) | 94.60 | 90.50 (-4%) | 95.07 | 95.89 (+1%) | 89.08 | 91.36 (+3%) |
| RF | 66.88 | 66.67 (0%) | 61.14 | 61.22 (+0%) | 87.01 | 86.54 (-1%) | 79.68 | 78.79 (-1%) |
| SVM | 94.42 | 86.61 (-8%) | 94.59 | 85.43 (-10%) | 97.24 | 94.65 (-3%) | 94.02 | 89.25 (-5%) |
| Mean | 72.28 | 77.61 (+7%) | 94.60 | 86.67 (-9%) | 85.97 | 85.80 (+0%) | 75.44 | 74.75 (-1%) |

**Supplementary Table S17. Cross-validation results when using the dataset obtained with random forests.**

| Performances | | | | | | | | |
|---|---|---|---|---|---|---|---|---|
| Model | Accuracy (AD) | Accuracy (AP) | Recall (AD) | Recall (AP) | Precision (AD) | Precision (AP) | Specificity (AD) | Specificity (AP) |
| AB | 59.19 | 66.25 (+12%) | 59.77 | 68.16 (+14%) | 75.97 | 79.97 (+5%) | 57.91 | 62.00 (+7%) |
| DT | 54.08 | 57.40 (+6%) | 54.90 | 58.07 (+6%) | 71.91 | 74.58 (+4%) | 52.25 | 55.92 (+7%) |
| GP | 37.25 | 69.02 (+85%) | 9.85 | 100.00 (+915%) | 92.66 | 69.02 (-26%) | 98.26 | 0.057 (-100%) |

| | | | | | | | | |
|---|---|---|---|---|---|---|---|---|
| KNN | 68.99 | 69.12 **(+0%)** | 99.98 | 99.94 **(+0%)** | 69.00 | 69.10 **(+0%)** | 0.00 | 0.49 **(+0%)** |
| MLP | 72.43 | 63.11 **(-13%)** | 82.87 | 65.76 **(-21%)** | 78.41 | 77.38 **(-1%)** | 49.19 | 57.21 **(+16%)** |
| GNB | 65.78 | 69.22 **(+5%)** | 59.21 | 61.50 **(+4%)** | 87.07 | 90.96 **(+4%)** | 80.42 | 86.40 **(+7%)** |
| RF | 52.35 | 57.90 **(+11%)** | 45.65 | 51.79 **(+13%)** | 75.63 | 80.19 **(+6%)** | 67.25 | 71.51 **(+6%)** |
| SVM | 51.93 | 68.22 **(+31%)** | 36.66 | 67.75 **(+85%)** | 85.29 | 83.06 **(-3%)** | 85.92 | 69.25 **(-19%)** |
| Mean | 56.64 | 67.24 **(+19%)** | 57.06 | 66.76 **(+17%)** | 77.19 | 78.68 **(+2%)** | 62.58 | 59.61 **(-5%)** |

Pairwise comparisons (ADs versus APs) between the metrics of each model for the different configurations are provided in Supplementary Table S18. The performance of APs was superior to that of ADs for all the metrics.

**Supplementary Table S18. Comparison of performance between datasets obtained with APs and ADs.**

| Performance | Accuracy | Recall | Precision | Specificity |
|---|---|---|---|---|
| AP > AD | 33 (69%) | 30 (63%) | 25 (52%) | 28 (58%) |
| AD > AP | 10 (21%) | 12 (25%) | 15 (31%) | 15 (31%) |
| AD = AP | 5 (10%) | 6 (12%) | 8 (17%) | 5 (11%) |

Another important step was to establish which FS techniques were appropriate for correctly predicting the different rust phenotypic groups. The superior boundaries for each confidence interval created for the mean values are shown in Supplementary Table S19. Accuracy was the only measure that did not have a significant p-value for the Shapiro-Wilk normality test (0.09678); thus, to quantify the confidence interval, we considered a normal distribution. For recall, precision and specificity, on the other hand, we used a Wilcoxon test to calculate these boundaries.

**Supplementary Table S19. Confidence intervals' superior boundaries calculated for the performance metrics.**

| | Accuracy | Recall | Precision | Specificity |
|---|---|---|---|---|
| 95% | 69.72 | 76.48 | 82.62 | 70.36 |
| 99% | 70.61 | 77.73 | 83.25 | 71.84 |
| 99.9% | 71.67 | 79.09 | 84.06 | 73.54 |

Using these values, we counted the quantity of results in each FS that exceeded these boundaries to check for significant improvements in performances. These counts are listed in Supplementary Tables S20-S23. Using different restrictions on the confidence intervals, we reached the same conclusions: FS1, FS2 and FS4 outperformed the other methods.

**Supplementary Table S20. Model accuracy performance and the confidence interval values.**

| | All | FS1 | FS2 | FS3 | FS4 | FS5 |
|---|---|---|---|---|---|---|
| 95% | 0 (0%) | 11 (69%) | 10 (63%) | 1 (6%) | 11 (69%) | 1 (6%) |
| 99% | 0 (0%) | 8 (50%) | 10 (63%) | 1 (6%) | 11 (69%) | 1 (6%) |
| 99.9% | 0 (0%) | 7 (44%) | 7 (44%) | 1 (6%) | 11 (69%) | 1 (6%) |

**Supplementary Table S21. Model recall performance and the confidence interval values.**

| | All | FS1 | FS2 | FS3 | FS4 | FS5 |
|---|---|---|---|---|---|---|
| 95% | 3 (19%) | 10 (63%) | 10 (63%) | 4 (25%) | 10 (63%) | 4 (25%) |
| 99% | 3 (19%) | 10 (63%) | 9 (56%) | 4 (25%) | 10 (63%) | 4 (25%) |
| 99.9% | 3 (19%) | 9 (56%) | 9 (56%) | 4 (25%) | 10 (63%) | 4 (25%) |

**Supplementary Table S22. Model precision performance and the confidence interval values.**

| | All | FS1 | FS2 | FS3 | FS4 | FS5 |
|---|---|---|---|---|---|---|
| 95% | 1 (6%) | 11 (69%) | 11 (69%) | 5 (31%) | 10 (63%) | 5 (31%) |
| 99% | 1 (6%) | 11 (69%) | 10 (63%) | 4 (25%) | 10 (63%) | 4 (25%) |
| 99.9% | 1 (6%) | 11 (69%) | 10 (63%) | 4 (25%) | 10 (63%) | 4 (25%) |

**Supplementary Table S23. Model specificity performance and the confidence interval values.**

| | All | FS1 | FS2 | FS3 | FS4 | FS5 |
|---|---|---|---|---|---|---|
| 95% | 2 (13%) | 9 (56%) | 9 (56%) | 4 (25%) | 10 (63%) | 5 (31%) |
| 99% | 2 (13%) | 9 (56%) | 9 (56%) | 4 (25%) | 8 (50%) | 4 (25%) |
| 99.9% | 1 (6%) | 7 (44%) | 9 (56%) | 4 (25%) | 8 (50%) | 4 (25%) |

The distribution of the different metrics for each FS can be visualized in Supplementary Figs. S9-S12, together with the groups identified by the Tukey test. The accuracy, recall and precision metrics were significantly different based on ANOVA. Supplementary Fig. S9 shows the group with the highest performance (FS1, FS2 and FS4). This same behavior is shown in the boxplots in Supplementary Figs. S10 and S11, with the exception that more groups were formed. However, considering the separation identified by the Tukey test of accuracy metrics, the recall and precision groups are divisions of the previous groups with a significant but small difference.

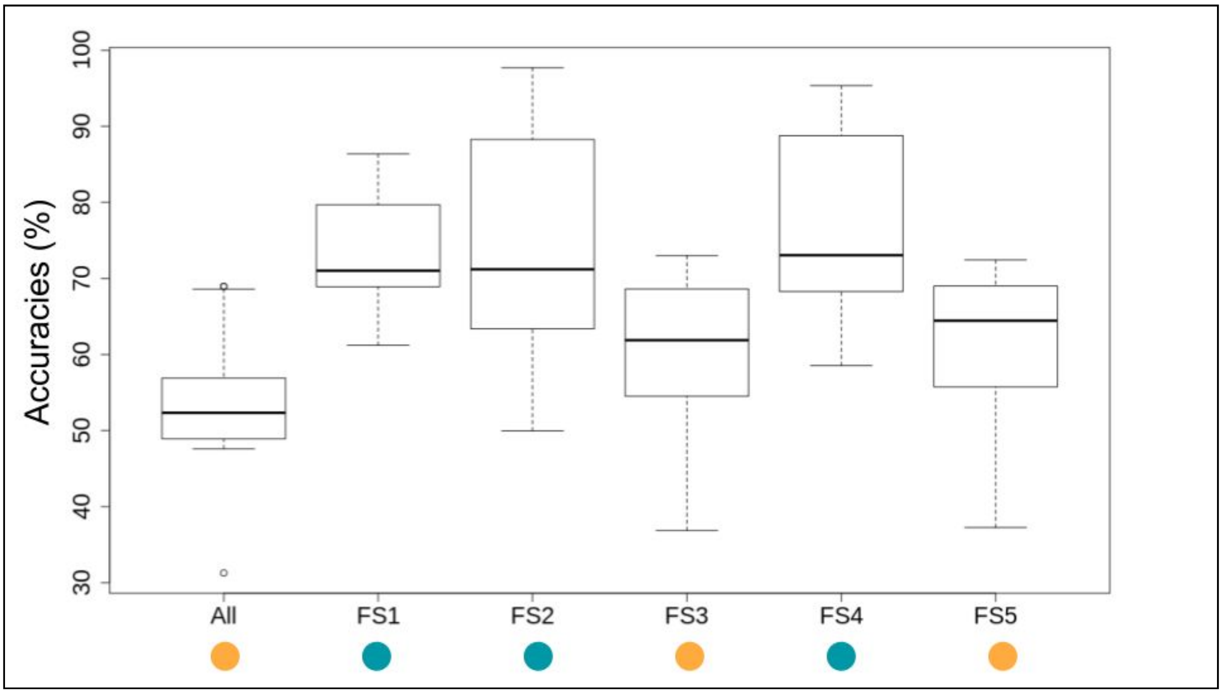


**Supplementary Fig. S9. Accuracy distribution for each subset created based on the different FS techniques. The colors are based on the groups identified with multiple comparisons by Tukey's test (ANOVA p-value of 0.000000000000455).**

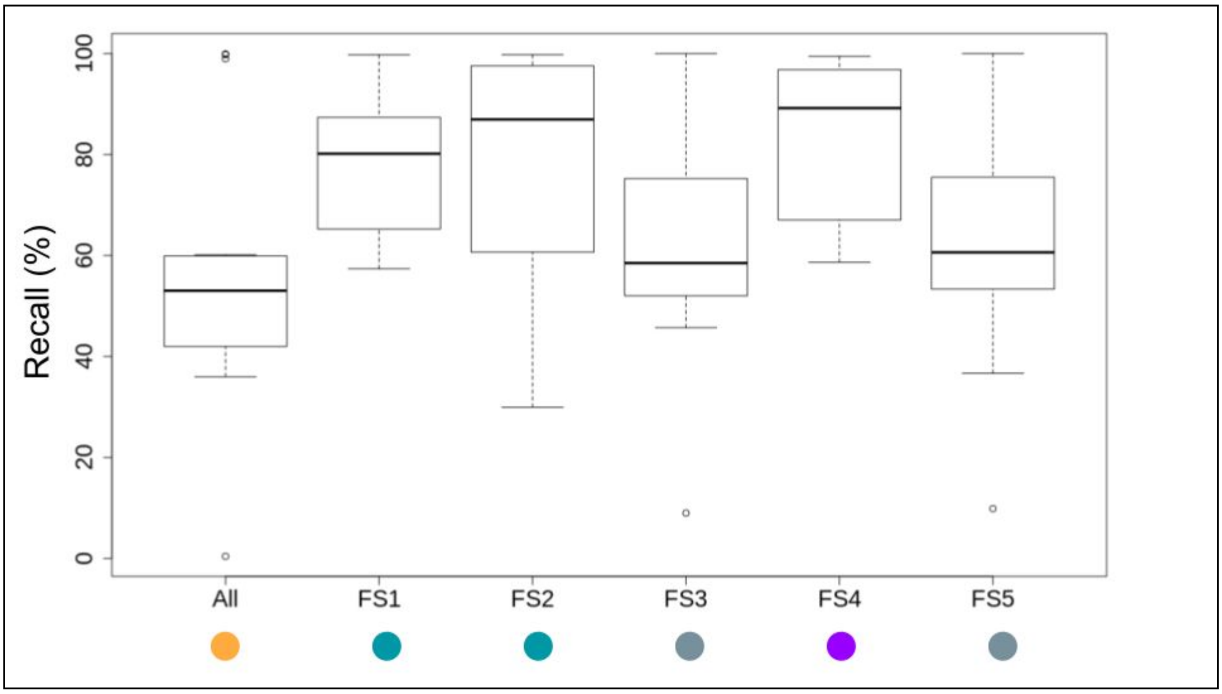


**Supplementary Fig. S10. Recall metric distribution for each subset created based on the different FS techniques. The colors are based on the groups identified with multiple comparisons by Tukey's test (ANOVA p-value of 0.000005375).**

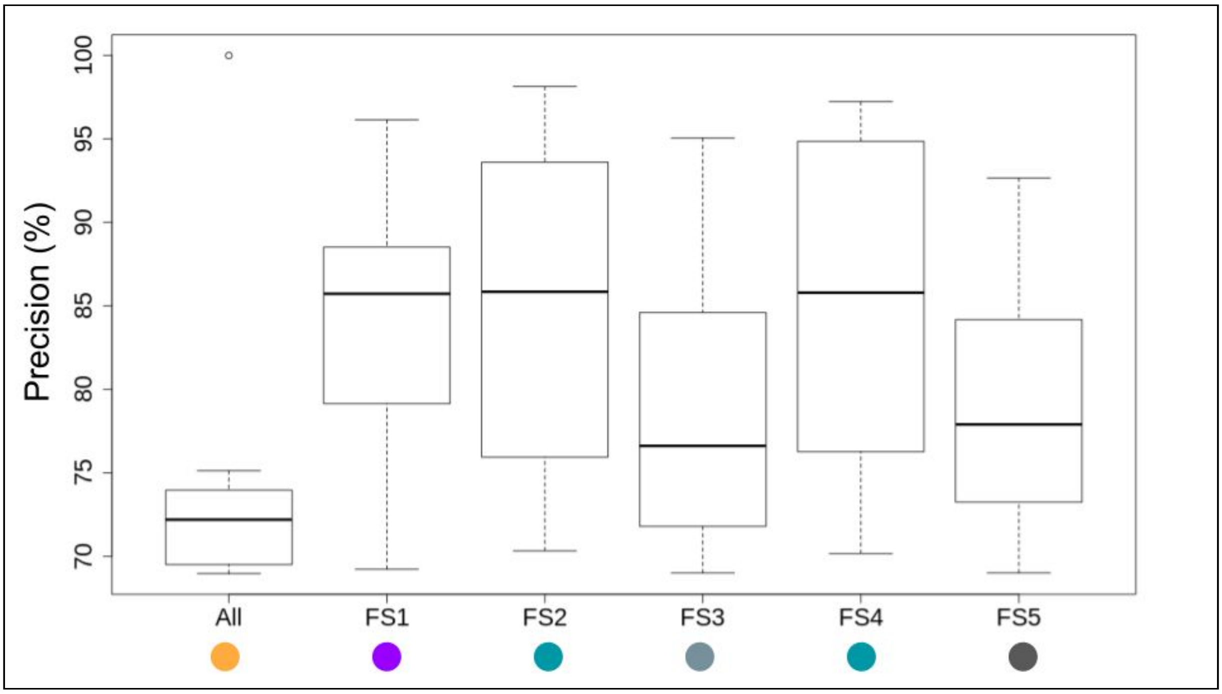


**Supplementary Fig. S11. Precision metric distribution for each subset created based on the different FS techniques. The colors are based on the groups identified with multiple comparisons by Tukey's test (ANOVA p-value of 0.000000673698).**

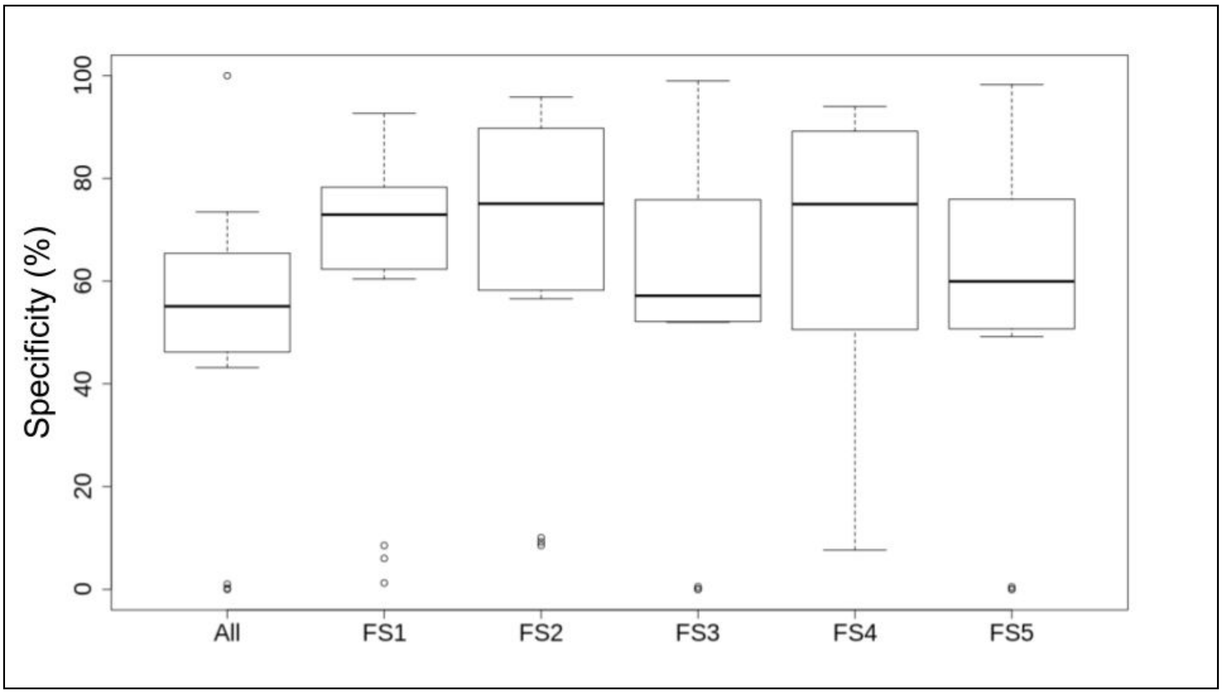


**Supplementary Fig. S12. Specificity metric distribution for each subset created based on the different FS techniques (ANOVA p-value of 0.0656).**

Considering FS1, FS2 and FS4 as the most promising strategies for selecting phenotype-associated variants, we selected SNPs identified with all of these methods (6 SNPs) to check for improvements in the evaluated metrics, as shown in Supplementary Table S24. Although individual FS techniques performed better than this dataset, we observed a substantial increase in these metrics when considering the initial dataset with 14,540 SNPs.

**Supplementary Table S24. Cross-validation results when using the SNPs selected by all of the different FS techniques.**

| Performances | | | | |
|---|---|---|---|---|
| Model | Accuracy (AD) | Recall (AD) | Precision (AD) | Specificity (AD) |
| AB | 73.06 | 72.92 | 85.90 | 73.36 |
| DT | 69.32 | 70.14 | 82.78 | 67.51 |
| GP | 72.57 | 72.92 | 85.20 | 71.79 |
| KNN | 70.13 | 73.55 | 81.38 | 62.53 |
| MLP | 71.98 | 72.62 | 84.60 | 70.57 |
| GNB | 73.32 | 71.53 | 87.52 | 77.28 |
| RF | 71.37 | 68.00 | 87.75 | 78.87 |
| SVM | 74.20 | 78.69 | 83.04 | 64.21 |

| | | | | |
|---|---|---|---|---|
| Mean | 72.28 | 72.77 | 84.90 | 71.18 |
| Mean FS1 | 77.80 | 80.54 | 86.89 | 72.94 |
| Mean FS2 | 82.01 | 89.60 | 86.70 | 76.69 |
| Mean FS4 | 77.61 | 86.67 | 85.80 | 74.75 |
| Mean All | 54.50 | 54.10 | 72.65 | 55.10 |

To evaluate the models' performance across the different datasets and check their appropriateness in predicting phenotypic groups, ROC curves were constructed, as shown in Supplementary Figs. S13-S20. The model predictive capability changed depending on the dataset configuration, as previously demonstrated.

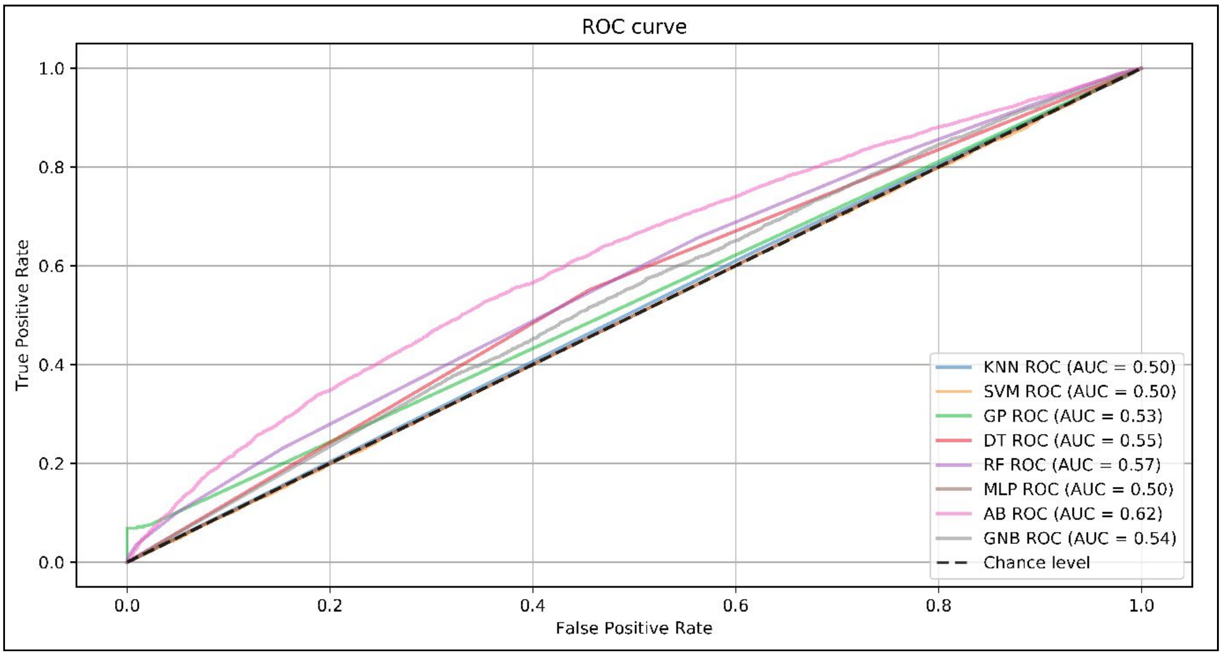


**Supplementary Fig. S13. ROC curves of model performance obtained using the entire dataset.**

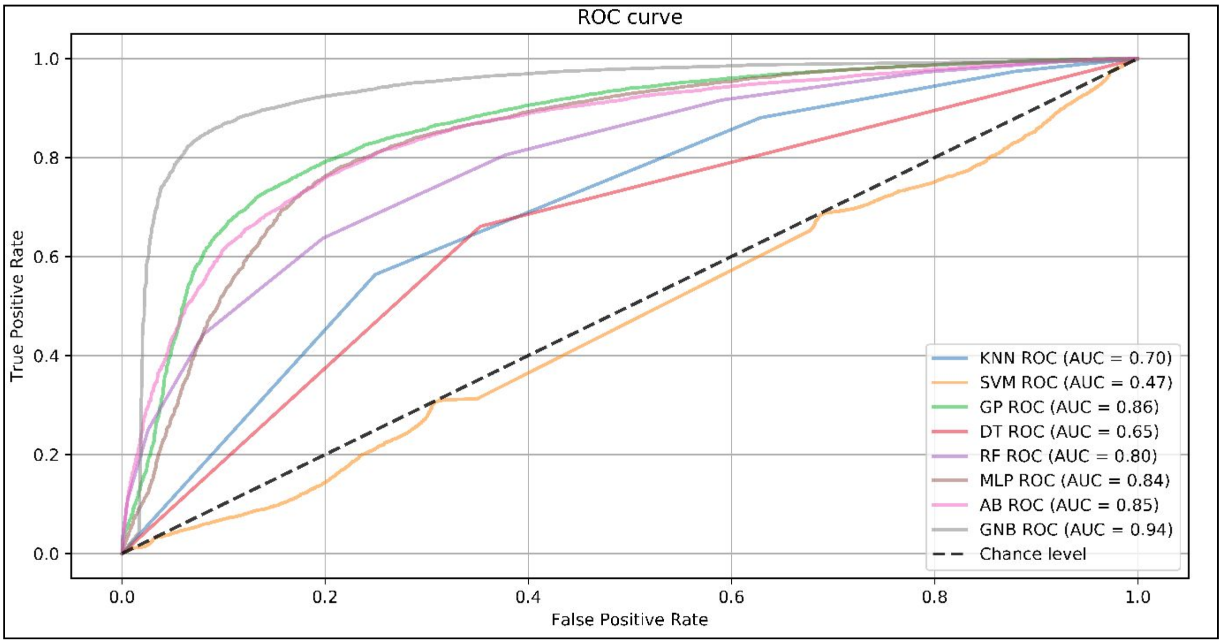


**Supplementary Fig. S14. ROC curves of model performance obtained using the dataset selected by gradient tree boosting.**

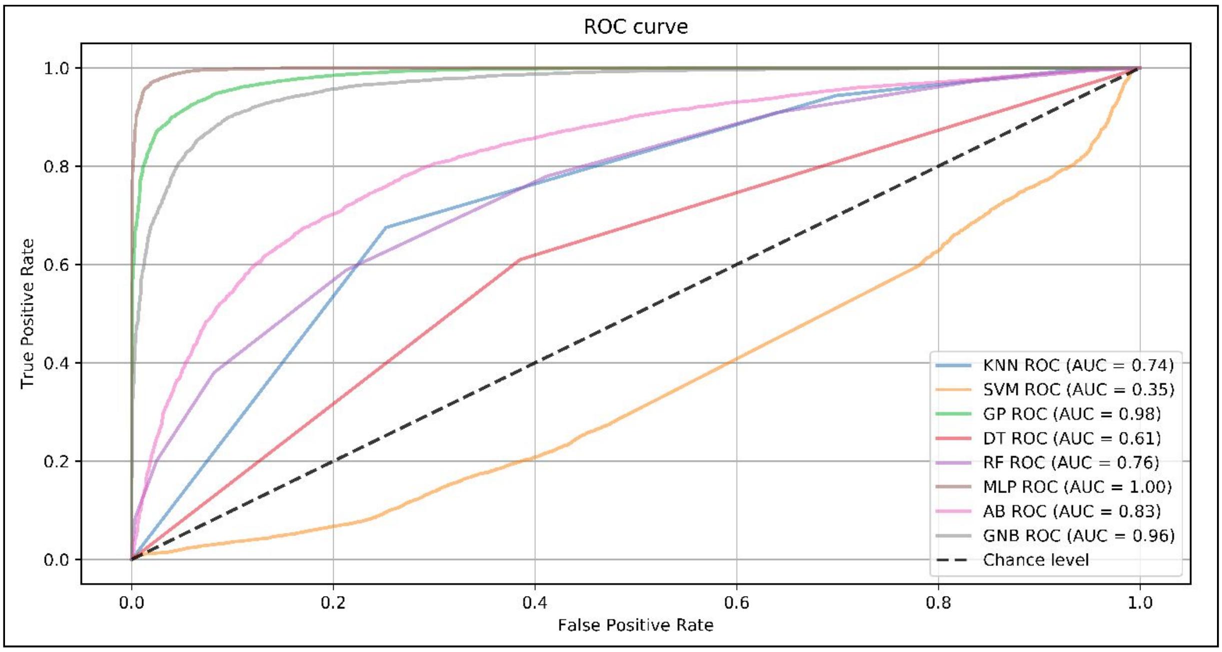


**Supplementary Fig. S15. ROC curves of model performance obtained using the dataset selected by L1-based SVC.**

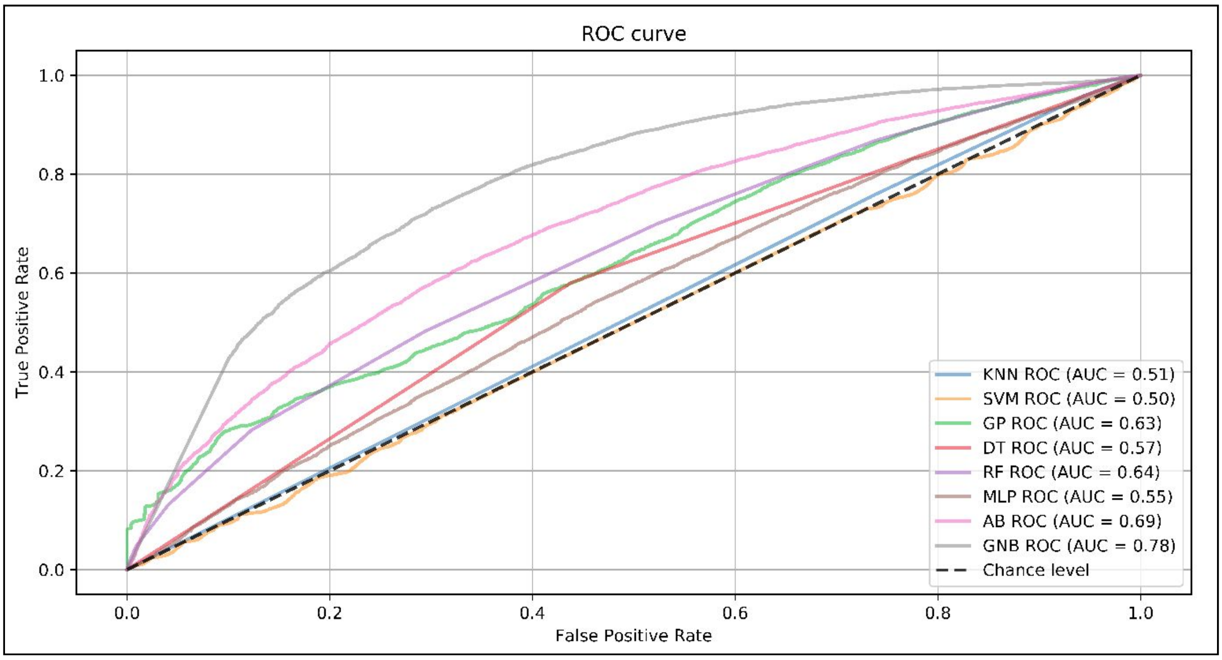


**Supplementary Fig. S16. ROC curves of model performance obtained using the dataset selected by extra trees.**

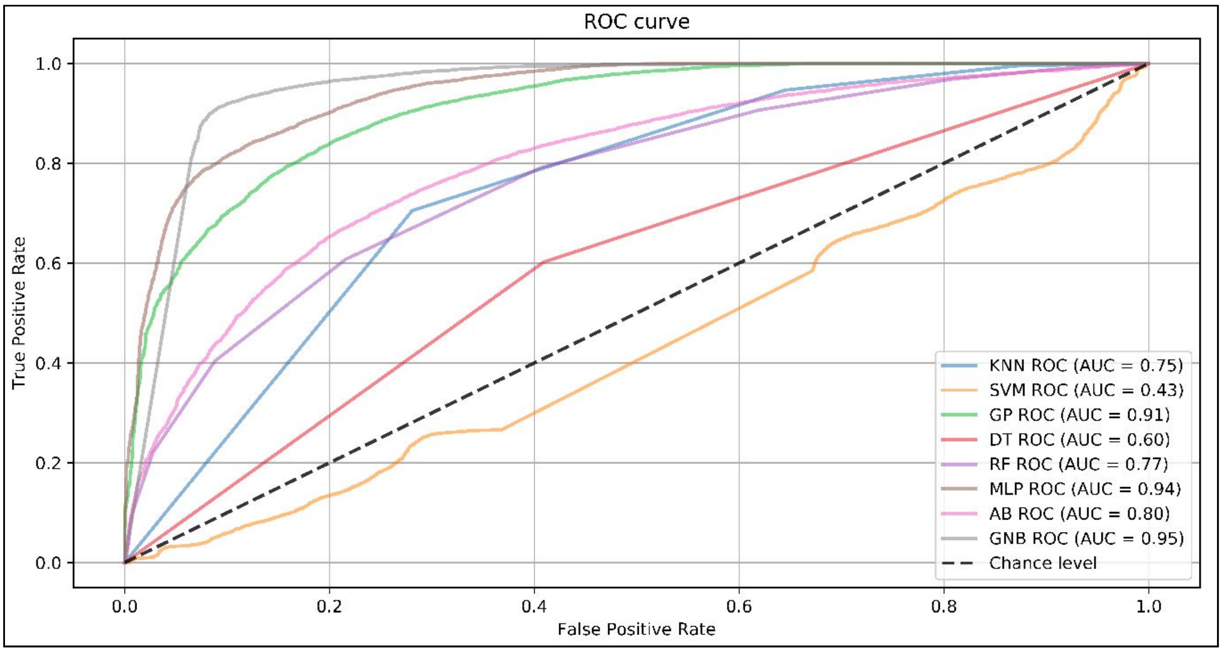


**Supplementary Fig. S17. ROC curves of model performance obtained using the dataset selected by the F-statistic from ANOVA.**

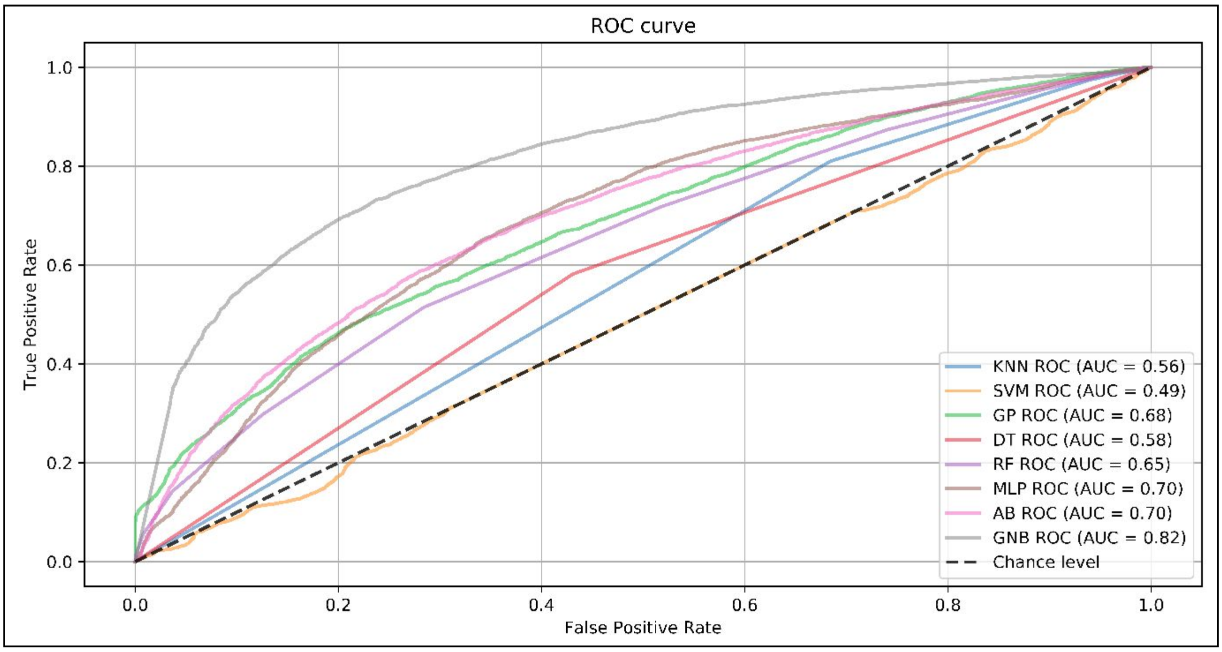


**Supplementary Fig. S18. ROC curves of model performance using the dataset obtained with random forests.**

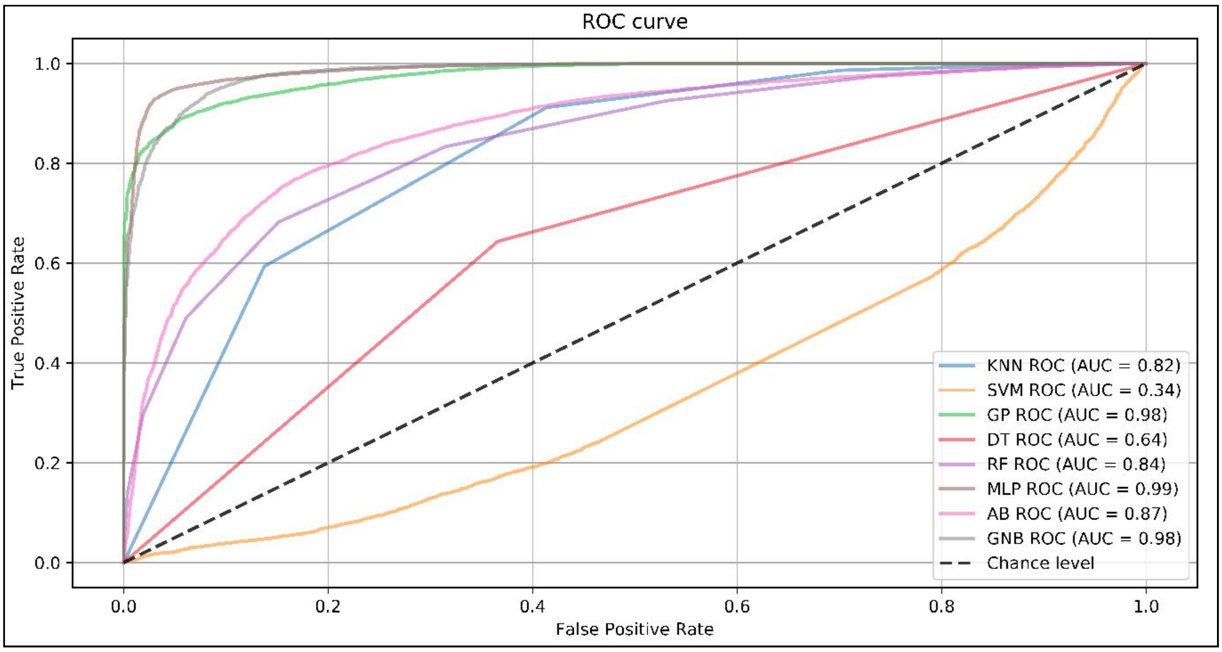


**Supplementary Fig. S19. ROC curves of model performance obtained using the dataset selected by the intersection of at least two of the datasets obtained using gradient tree boosting, L1-based SVC or the F statistic from ANOVA.**

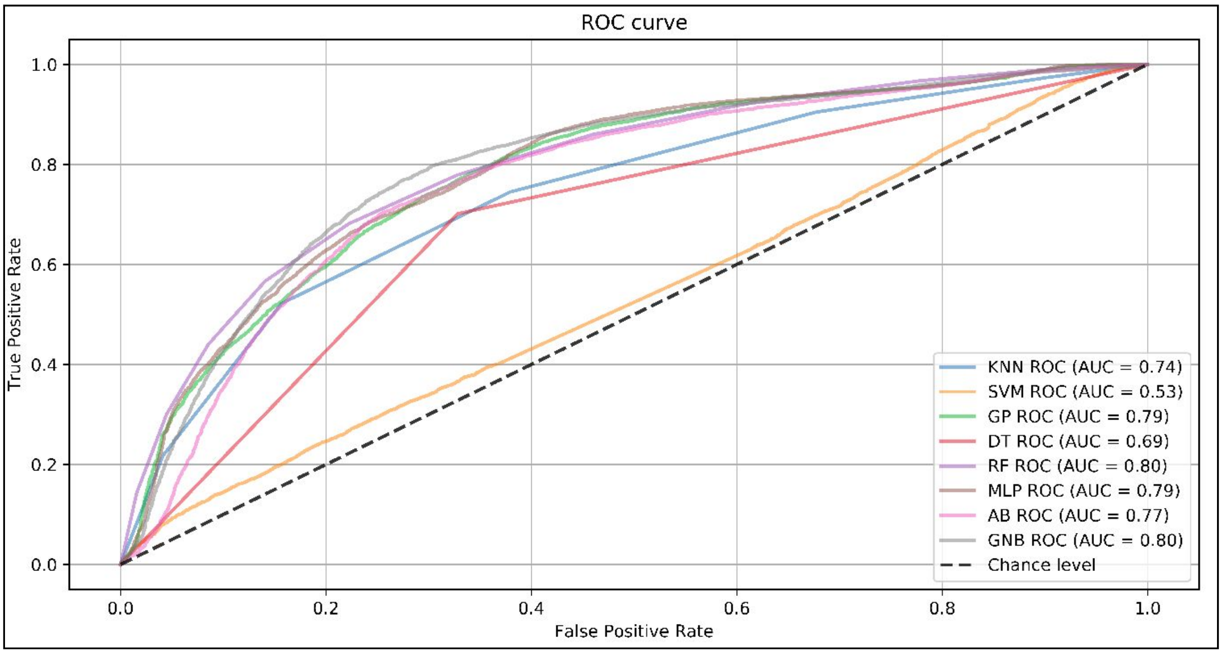


**Supplementary Fig. S20. ROC curves of model performance obtained using the dataset selected by the intersection of the datasets obtained using gradient tree boosting, L1-based SVC and the F statistic from ANOVA.**

According to the ROC curves, the best ML strategies for predicting the phenotypic groups were MLP, GNB and GP. The ROC curves based on these algorithms and the different dataset configurations can be visualized in Supplementary Figs. S21-S23. The best performances were observed for FS2 and the intersection of two FSs (I2).

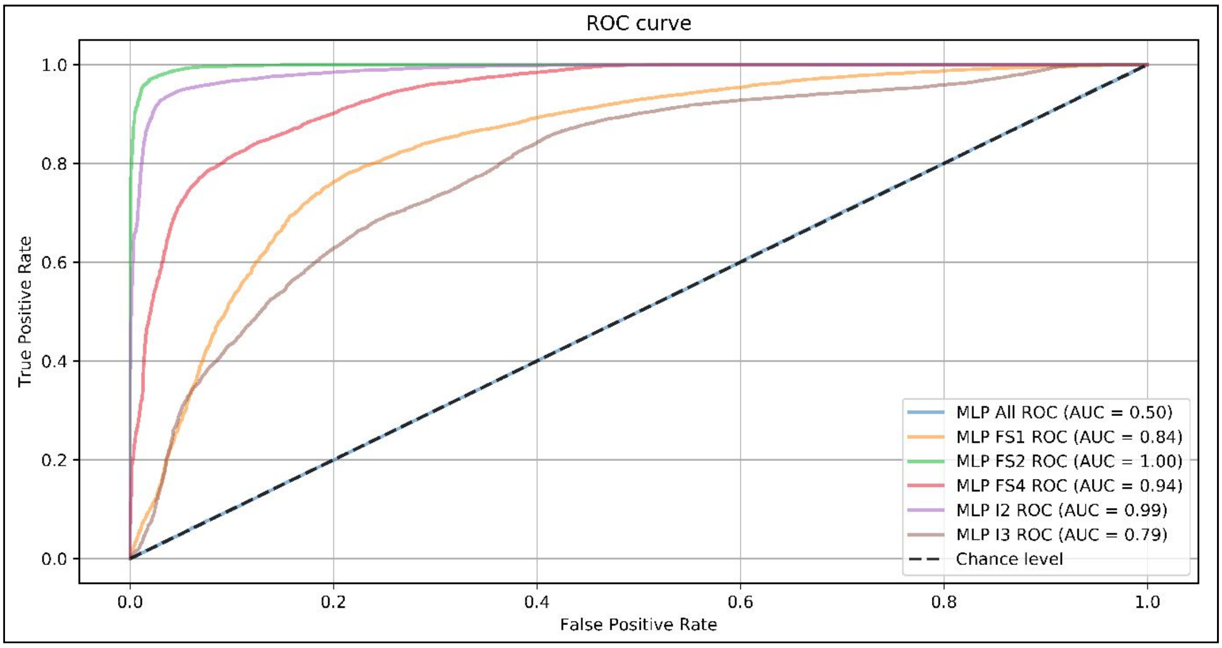


**Supplementary Fig. S21. MLP results and different dataset configurations.**

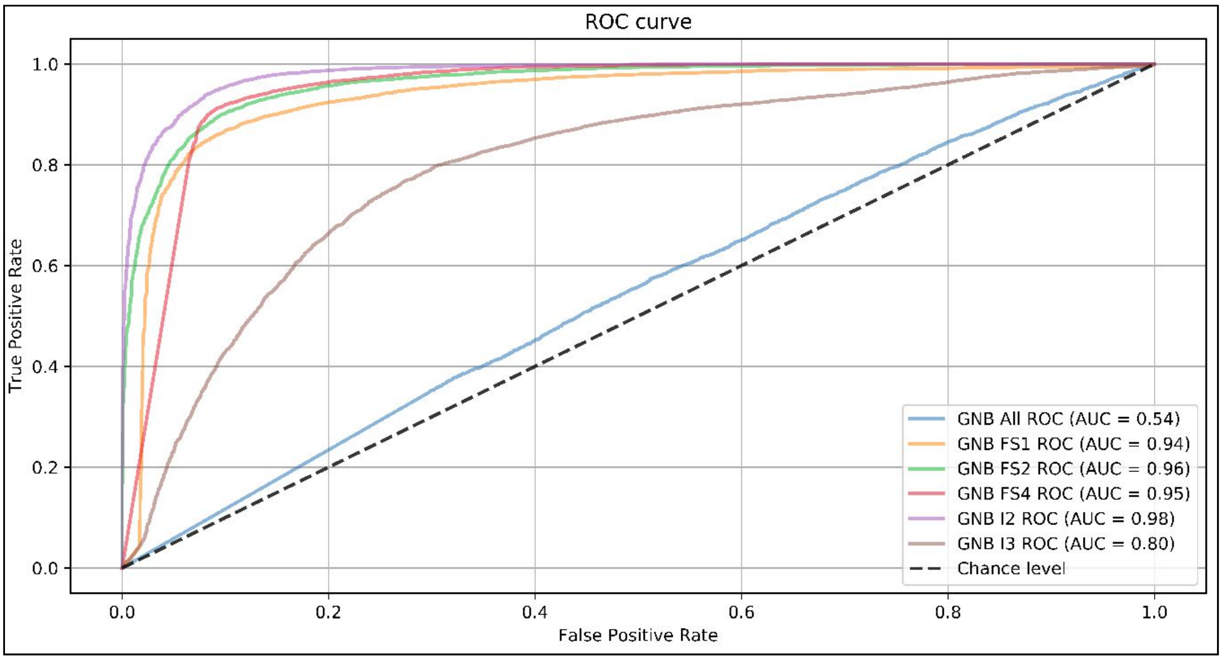


**Supplementary Fig. S22. GNB results and different dataset configurations.**

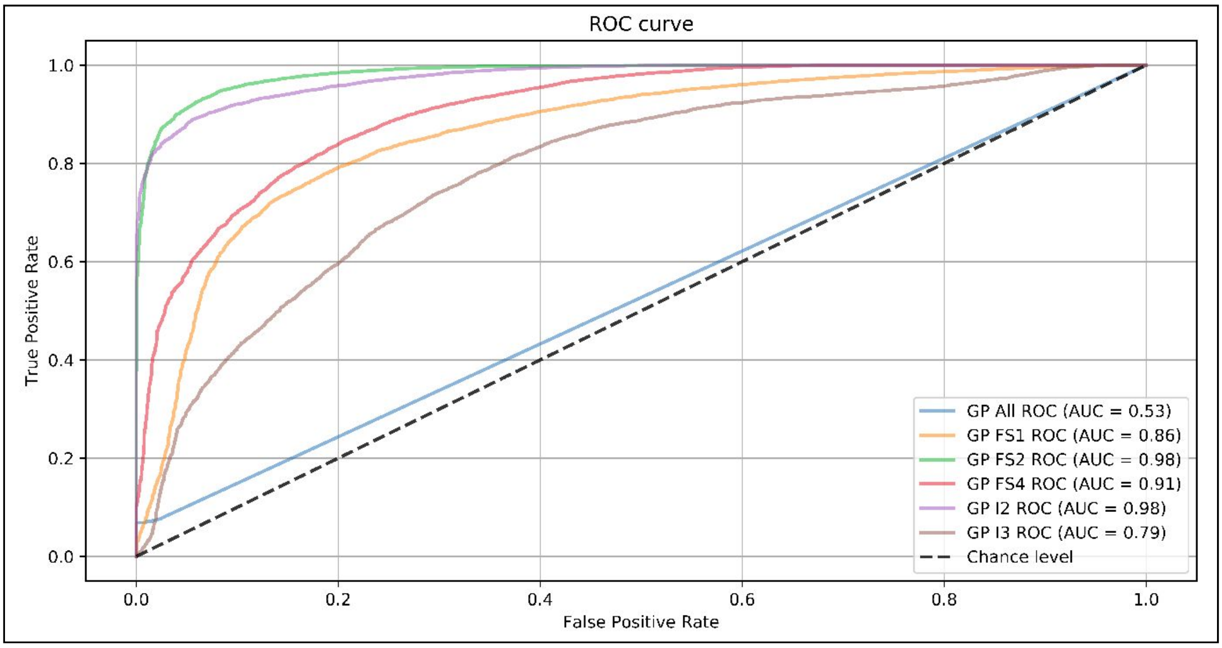


**Supplementary Fig. S23. GP results and different dataset configurations.**

Using sets of *S. spontaneum* chromosomes with allele specificity, we plotted a physical representation of the organization of subsets of CDSs related to BACs and Inter 2 SNPs (Supplementary Fig. S24). These alignment lengths covered at least 75% of the sequence of BACs or scaffolds containing Inter 2 SNPs with a high similarity established, showing the elevated probability of correspondences. The mean alignment length between BACs and CDSs was 1,350 (ranging from 80 to 13,466), and that between scaffolds and CDSs was 285 (ranging from 78 to 4,634).

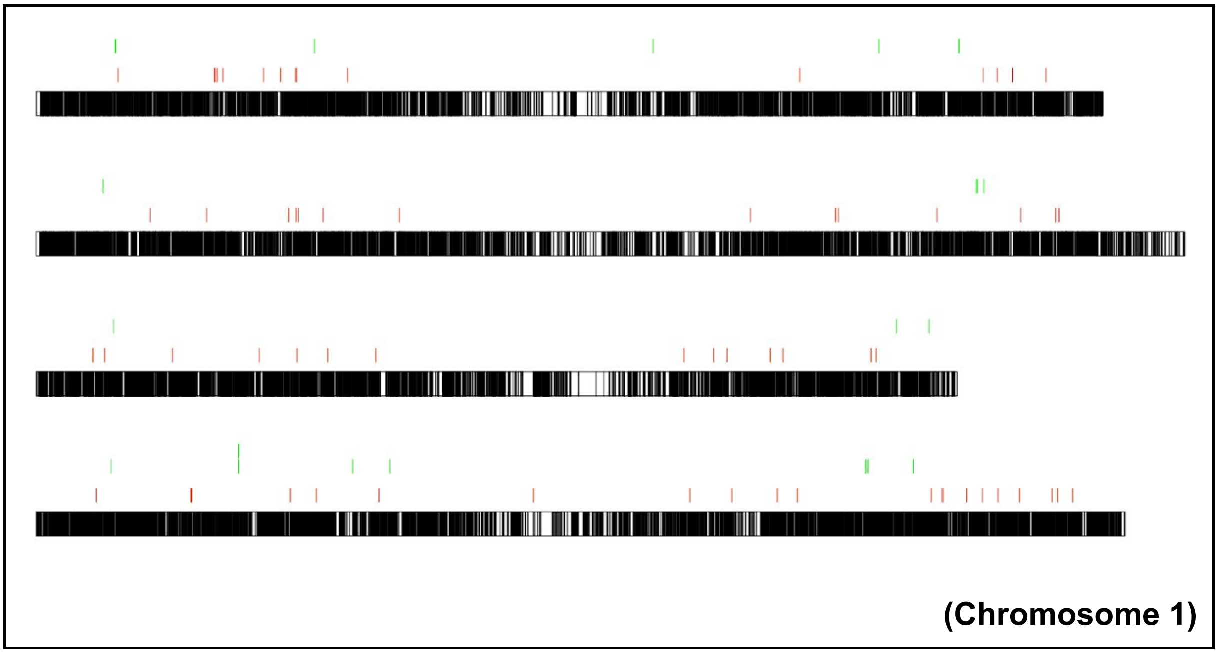

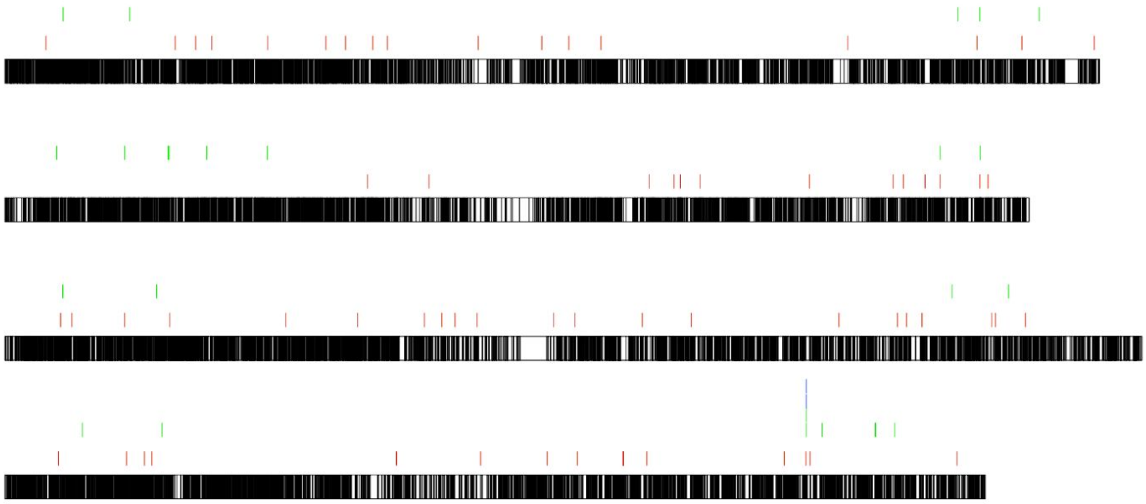

**(Chromosome 2)**

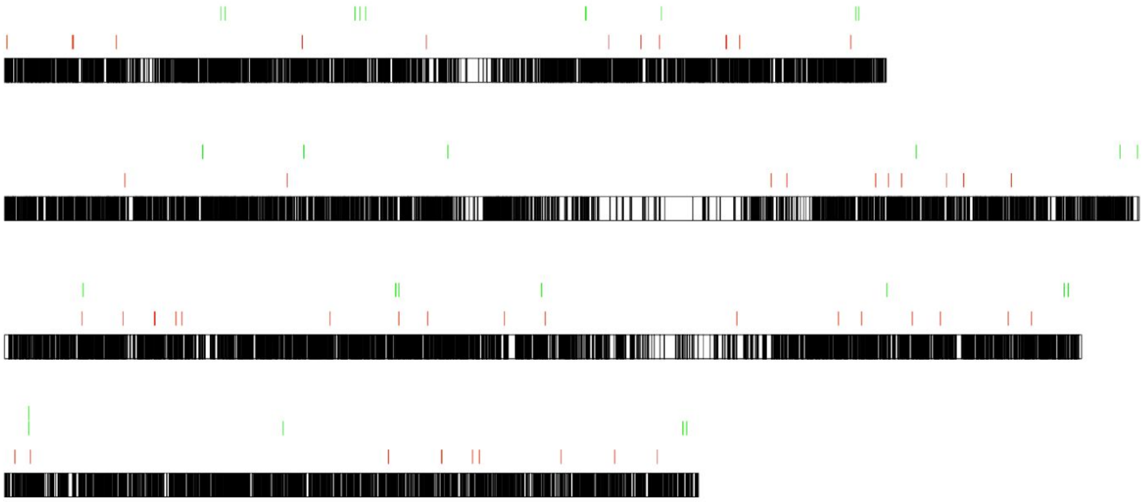

**(Chromosome 3)**

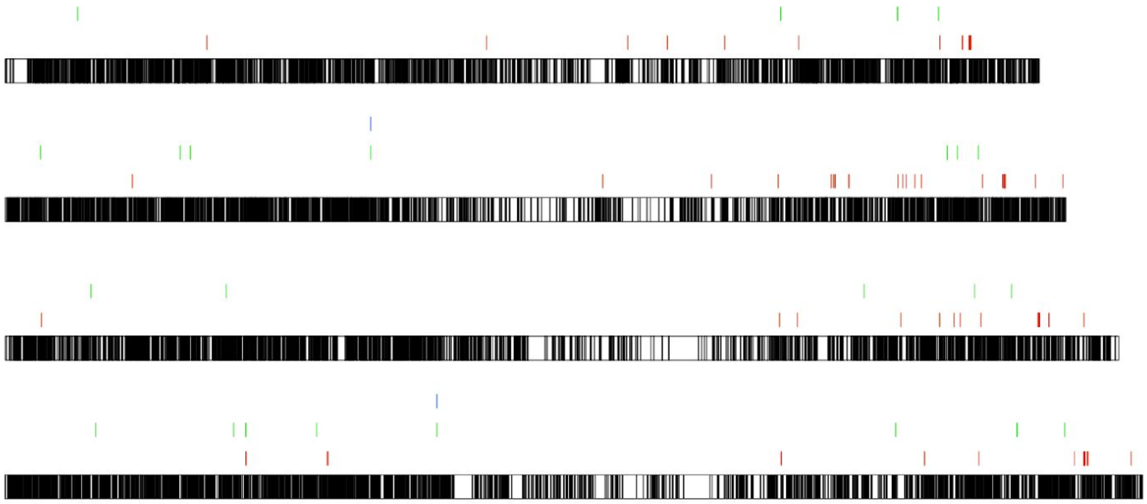

**(Chromosome 4)**

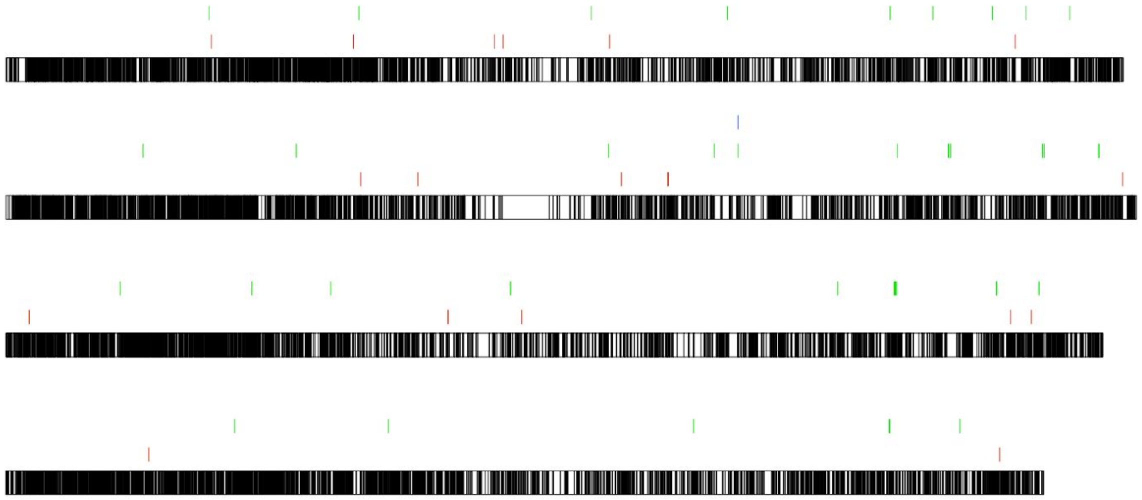

(Chromosome 5)

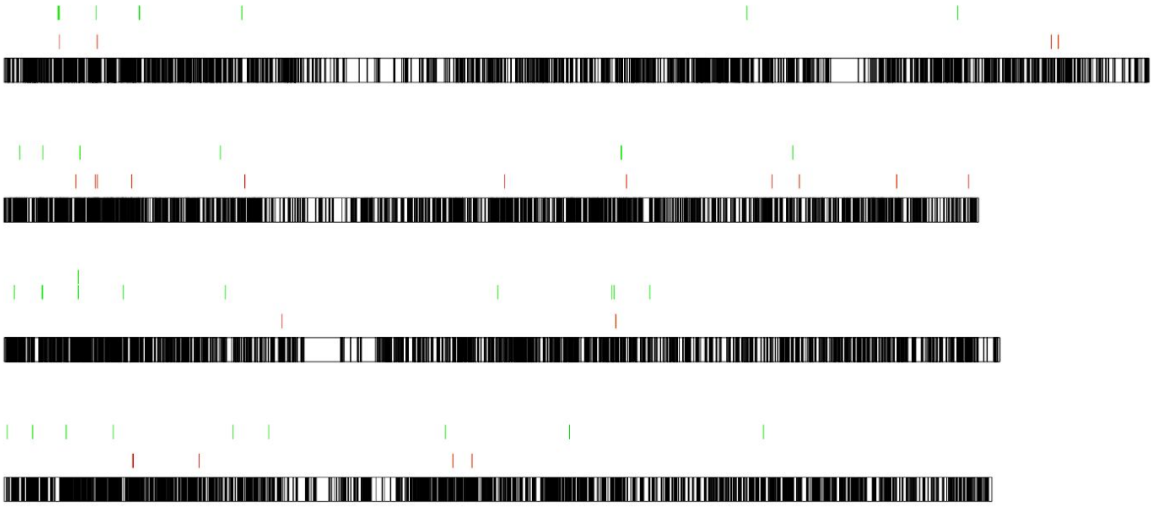

(Chromosome 6)

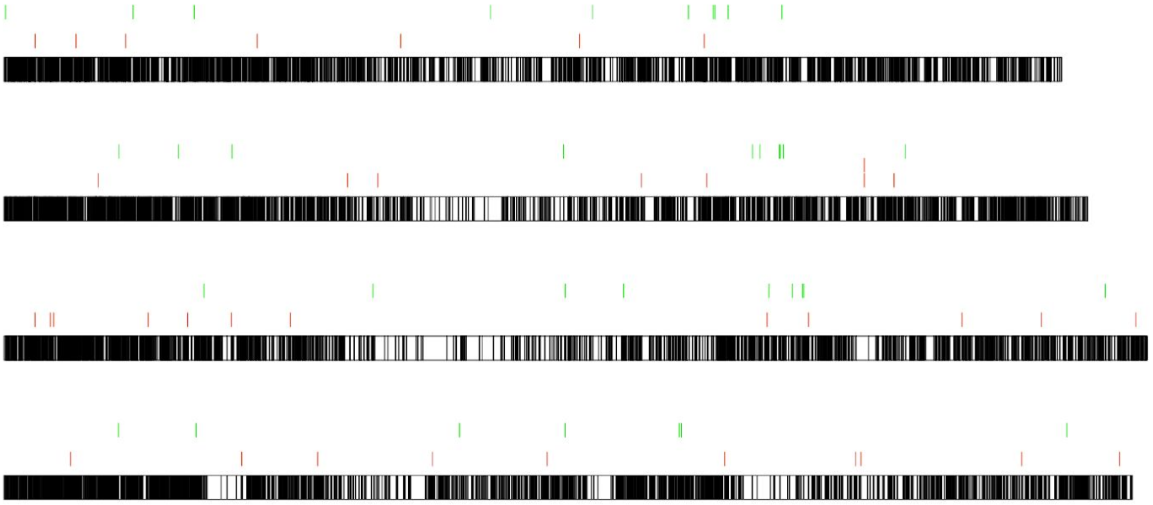

(Chromosome 7)

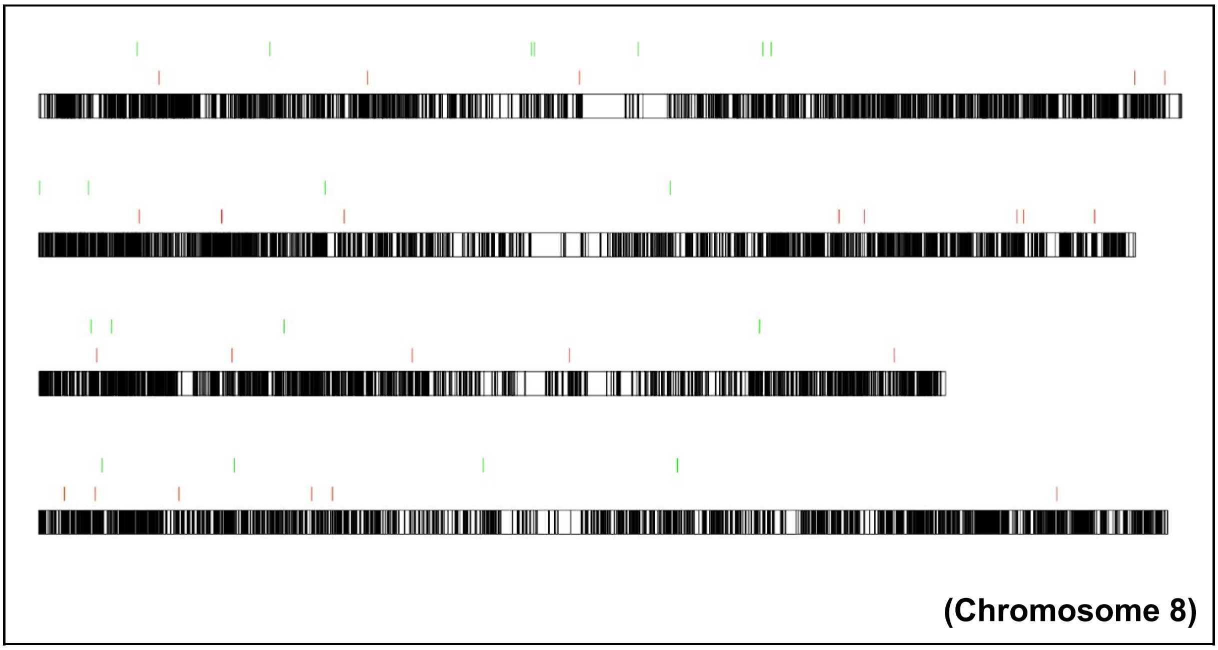


**Supplementary Fig. S24. Representation of CDS distributions across *S. spontaneum* chromosomes. Each chromosome is shown in a different block with the four alleles. The karyotype is colored according to the presence of CDSs (black bands). The locations of CDSs corresponding to brown rust BACs are colored in red, and the locations of CDSs corresponding to scaffolds containing the selected SNPs are colored in green.**

There were different *S. spontaneum* CDSs related to (I) BACs and (II) Inter 2 scaffolds, as shown in Supplementary Fig. S24. From these CDSs, it was possible to extract 50 GO categories shared by these two groups and 50 exclusive to (II). We also performed enrichment analysis of these 50 exclusive GO categories, the results of which are shown in Supplementary Fig. S25.

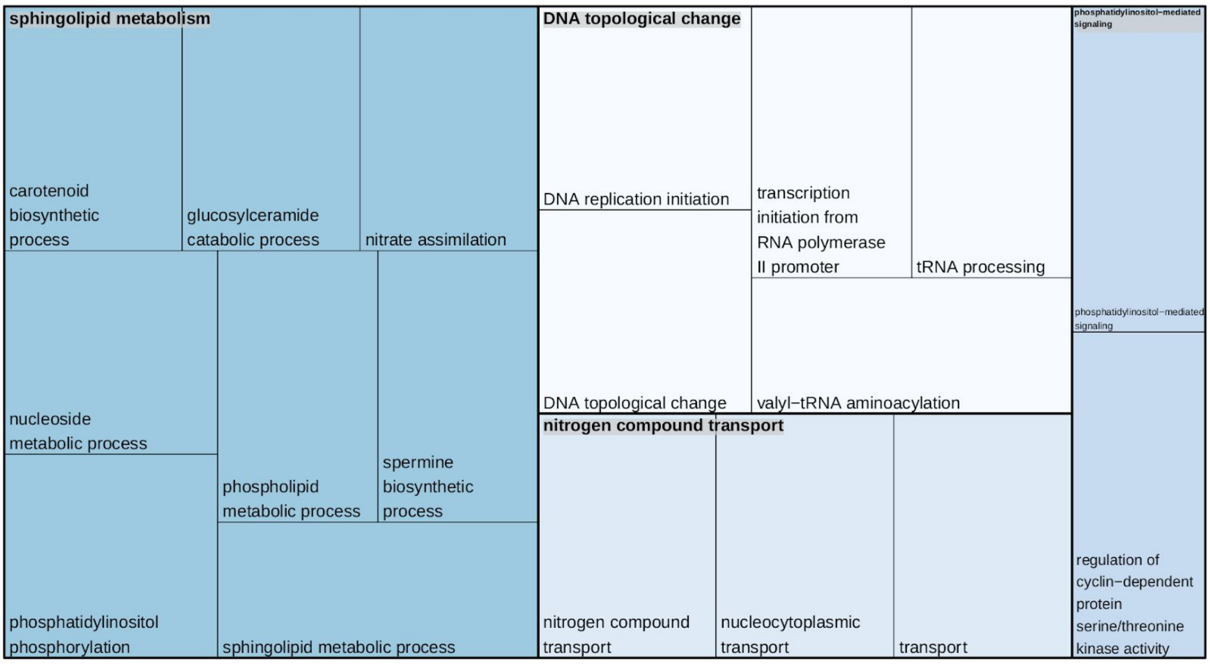


**Supplementary Fig. S25. GO categories found in the CDSs corresponding to scaffolds containing the selected SNPs and not in the BACs' correspondences.**

In addition to GO categories, we also identified corresponding metabolic pathways in (I) and (II), as follows:

**Pathways related to CDSs corresponding to BACs:**

1. path:sbi00052 - Galactose metabolism
2. path:sbi01230 - Biosynthesis of amino acids
3. path:sbi03013 - RNA transport
4. path:sbi00280 - Valine, leucine and isoleucine degradation
5. path:sbi04144 - Endocytosis
6. path:sbi03040 - Spliceosome
7. path:sbi00561 - Glycerolipid metabolism
8. path:sbi00790 - Folate biosynthesis
9. path:sbi00900 - Terpenoid backbone biosynthesis
10. path:sbi00240 - Pyrimidine metabolism
11. path:sbi04141 - Protein processing in endoplasmic reticulum
12. path:sbi03010 - Ribosome
13. path:sbi00040 - Pentose and glucuronate interconversions
14. path:sbi01212 - Fatty acid metabolism
15. path:sbi03008 - Ribosome biogenesis in eukaryotes
16. path:sbi00190 - Oxidative phosphorylation
17. path:sbi00564 - Glycerophospholipid metabolism
18. path:sbi00860 - Porphyrin and chlorophyll metabolism
19. path:sbi04070 - Phosphatidylinositol signaling system
20. path:sbi04145 - Phagosome
21. path:sbi00500 - Starch and sucrose metabolism
22. path:sbi00250 - Alanine, aspartate and glutamate metabolism

23. path:sbi03020 - RNA polymerase
24. path:sbi00960 - Tropane, piperidine and pyridine alkaloid biosynthesis
25. path:sbi01100 - Metabolic pathways
26. path:sbi00400 - Phenylalanine, tyrosine and tryptophan biosynthesis
27. path:sbi00051 - Fructose and mannose metabolism
28. path:sbi04016 - MAPK signaling pathway - plant
29. path:sbi00780 - Biotin metabolism
30. path:sbi04146 - Peroxisome
31. path:sbi00061 - Fatty acid biosynthesis
32. path:sbi01110 - Biosynthesis of secondary metabolites
33. path:sbi00970 - Aminoacyl-tRNA biosynthesis
34. path:sbi03440 - Homologous recombination
35. path:sbi03018 - RNA degradation
36. path:sbi00230 - Purine metabolism
37. path:sbi00591 - Linoleic acid metabolism
38. path:sbi00270 - Cysteine and methionine metabolism
39. path:sbi00260 - Glycine, serine and threonine metabolism
40. path:sbi00010 - Glycolysis / Gluconeogenesis
41. path:sbi04626 - Plant-pathogen interaction

**Pathways related to CDSs corresponding to scaffolds found in the intersection analysis:**

1. path:sbi03010 - Ribosome
2. path:sbi00902 - Monoterpenoid biosynthesis
3. path:sbi00051 - Fructose and mannose metabolism
4. path:sbi00052 - Galactose metabolism
5. path:sbi00940 - Phenylpropanoid biosynthesis
6. path:sbi01230 - Biosynthesis of amino acids
7. path:sbi00030 - Pentose phosphate pathway
8. path:sbi00920 - Sulfur metabolism
9. path:sbi03013 - RNA transport
10. path:sbi00511 - Other glycan degradation
11. path:sbi00062 - Fatty acid elongation
12. path:sbi03022 - Basal transcription factors
13. path:sbi01110 - Biosynthesis of secondary metabolites
14. path:sbi00250 - Alanine, aspartate and glutamate metabolism
15. path:sbi00500 - Starch and sucrose metabolism
16. path:sbi00564 - Glycerophospholipid metabolism
17. path:sbi03040 - Spliceosome
18. path:sbi00970 - Aminoacyl-tRNA biosynthesis
19. path:sbi04120 - Ubiquitin mediated proteolysis
20. path:sbi03018 - RNA degradation
21. path:sbi00513 - Various types of N-glycan biosynthesis
22. path:sbi00230 - Purine metabolism

23. path:sbi00380 - Tryptophan metabolism
24. path:sbi00600 - Sphingolipid metabolism
25. path:sbi01200 - Carbon metabolism
26. path:sbi00510 - N-Glycan biosynthesis
27. path:sbi00010 - Glycolysis / Gluconeogenesis
28. path:sbi01100 - Metabolic pathways
29. path:sbi04626 - Plant-pathogen interaction
